# Cognitive and molecular characterization of the Ts66Yah murine model of Down syndrome: deepening on hippocampal changes associated with genotype and aging

**DOI:** 10.1101/2024.01.02.573811

**Authors:** Chiara Lanzillotta, Monika Rataj Baniowska, Francesca Prestia, Chiara Sette, Valérie Nalesso, Marzia Perluigi, Eugenio Barone, Arnaud Duchon, Antonella Tramutola, Yann Herault, Fabio Di Domenico

## Abstract

Down syndrome (DS) is the most common condition with intellectual disability and is caused by trisomy of *Homo sapiens* chromosome 21 (HSA21). The increased dosage of genes on HSA21 is the cause for the initial neurodevelopmental disorder and for further development of cognitive decline, however the molecular mechanisms promoting brain pathology along ageing are still missing. One of the major challenges in the study of DS is the lack of reliable murine model able to accurately replicate genotypic and phenotypic aspects observed in humans along ageing. Preclinical studies in DS were pioneered using the Ts65Dn murine model, which despite its genetic limitations, has been extremely helpful in characterising the progression of brain degeneration. The novel Ts66Yah model represents an evolution of the Ts65Dn, with phenotypes only induced by trisomic HSA21 homologous genes, closer to human DS condition. In this study, we confirmed the behavioural features of Ts66Yah mice with improvement in the detection of spatial memory defects and also a new anxiety-related phenotype. The molecular characterisation of Ts66Yah demonstrated the aberrant regulation of redox balance, proteostasis, stress response, metabolic pathways, programmed cell death and synaptic plasticity. Intriguingly, the genotype-related alterations of those pathways occur early promoting the alteration of brain development and the onset of a condition of premature aging. Overall, data collected in Ts66Yah provide novel and consolidated insights, devoid of genome bias, concerning trisomy-driven processes that contribute to brain pathology in conjunction with aging. This, in turn, aids in bridging the existing gap in comprehending the intricate nature of DS phenotypes.

## INTRODUCTION

Down syndrome (DS) is the most common genomic disorder of intellectual disability and is caused by trisomy of *Homo sapiens* chromosome 21 (HSA21). The DS phenotype involves manifestations that affect multiple bodily systems including the musculoskeletal, neurological and cardiovascular systems. The aging process is accelerated in people with DS compared with the general population, resulting in increased incidence of a range of medical conditions, among which the early onset Alzheimer’s disease (AD) [1]. The increased dosage of *APP* in trisomy 21 is necessary for increased AD risk in individuals with DS, although the underlying mechanisms that link *APP* dosage to neurodegeneration are unknown. Indeed, other genes, such as DYRK1A and CBS, are strongly linked to the clinical outcomes of DS neuropathology [2–5]. Nonetheless, a significant challenge for researchers studying Down syndrome lies in the identification of a mouse model with robust genetic validity that accurately manifests human DS-related phenotypes. This is essential for deciphering pertinent pathological mechanisms.

In the mouse genes orthologues to HSA21 are located in three syntenic regions on MMU10, MMU16 and MMU17 with the largest being on MMU16. To date, >20 mouse models that are trisomic for different HSA21 homologous portions of MMU16, MMU17 and MMU10 have been created. Noteworthy, the majority of data acquired in DS using mouse models were pioneered and strongly driven by the use of the Ts(17^16^)65Dn/J (Ts65Dn) mouse line. This model developed by Muriel Davidson and Roger Reeves [6, 7], carries around 100 protein coding genes orthologues to HSA21 in three copies on a supernumerary hybrid chromosome comprised of the centromere of MMU17 and from *Mrpl39* to the distal part of MMU16, orthologous to HSA21 [3]. However, the Ts65Dn mouse also carries three copies of the Mmu17-derived region, which includes proteins coding genes that are not orthologous to genes on HSA21. Thus, some of the gene-dosage effects seen in this strain may not relate to human DS [8, 9].

Mouse models of DS exhibit aging dependent changes to behaviour and cognition [3]. Most notably, the Ts65Dn and related models exhibits progressive cognitive decline [10, 11], which has been linked to early APP overexpression and subsequent increase of Aβ oligomers in the brain [11–14]. In addition, HSA21encoded DYRK1A has effects on the phosphorylation of TAU at T212 (associated with AD) and of mRNA splicing factors of the TAU encoding gene *MAPT*, which may promote the formation of neurofibrillary tangles (NFTs). Studies from our laboratory confirmed that the Ts65Dn and genetically similar models (Ts2Cje) demonstrate the alteration of redox balance, protein homeostasis (proteostasis) and energy metabolic pathways, including insulin signalling and mitochondrial processes, as observed in human DS. Furthermore, we observed that the molecular changes detected in these DS mice were reliant on genotype but worsened with aging, being directly or indirectly implicated in the development of neurodegenerative signatures and in the reduction of cognitive performances [15–17]. Data collected so far highlight the importance of murine models for the study of brain changes associated with DS phenotype and for identification of the molecular mechanisms promoting age-related neurodegeneration and will allow to propose in the near future additional druggable molecular targets. Nevertheless, pharmacological strategies aimed to rescue the DS phenotypes in mice has yielded several promising results that so far have not been fully replicated in human clinical trials [18, 19]. Such misalignment in translating pre-clinical into clinical studies might be associated, with some degree of confidence, with the paucity of DS mouse models. Indeed, the triplication of non-HSA21 orthologous genes may have relevant effects on mice phenotype since the neuro-developmental stage.

Recently, Herault and colleagues developed a new line named Ts66Yah, derived from the Ts65Dn lineage but no longer carrying the duplicated centromeric part of Mmu17 [20]. The Ts66Yah mouse model was generated using CrispR/Cas9 technology in vivo to remove the segment of DNA corresponding to the centromeric Mmu17 located on the Ts65Dn mini chromosome. Although this remains a partial DS model since the triplicated *Mrpl39 to Zbtb21* region encompasses 102/187 of the HSA21 orthologous protein-coding genes, it is certainly more genetically valid for DS studies than the Ts65Dn, while the general transmission of the recombined Ts66Yah chromosome remains similar to the Ts65Dn. Importantly, Ts66Yah and Ts65Dn mice were found to replicate similar behavioural alterations, characteristic of DS pathology, but Ts66Yah demonstrated a similar cognitive phenotype with no hyperactivity behaviour [20]. Delays in motor development, communication, and olfactory spatial memory were present in Ts66Yah but more pronounced in Ts65Dn neonates. Adult Ts66Yah mice showed working memory deficits and sex-specific effects in exploratory behaviour and spatial hippocampal memory, while long-term spatial memory was preserved [20]. Similarities for brain and craniofacial changes were observed in Ts66Yah and Ts65Dn. Interestingly, while the expression analysis in the hippocampus showed common dysregulation of main pathways, the entorhinal cortex (EC) results more affected in the Ts65Dn compared to Ts66Yah [20]. Further, many genes from the Mmu17 non-HSA21 orthologous region were overexpressed both in the hippocampus and the EC of the Ts65Dn. In a second report from Bianchi’s laboratory, authors demonstrated that several Mmu17 non-HSA21 orthologous genes are uniquely overexpressed in Ts65Dn embryonic forebrain, producing major differences in dysregulated genes and pathways [21]. Altogether the results from Ts65Dn and Ts66Yah mice highlighted the interference of the non-HSA21 orthologous region in the Ts65Dn, which, if under increased genetic dosage, may change penetrance or expressivity of features in individual with DS. Thus, the Ts66Yah model seems to provide a refined analysis of DS phenotypes during lifetime. Furthermore, Ts66Yah mice demonstrated a stronger construct validity for mimicking specific consequences of DS genetic overdosage, certainly more translatable to humans and with potential better proof-of-concept studies for therapeutics.

In this study, we conducted a comprehensive long-term behavioural analysis in the Ts66Yah model and examined age-dependent alterations in various molecular mechanisms and pathways associated with the neurodegenerative process. Our objective is to establish and validate the utility of Ts66Yah mice in investigating the molecular aspects of brain pathology linked to aging. We aim to consider both discrepancies and commonalities in respect of what has been observed in other murine models of DS.

## MATERIALS AND METHODS

### Animals

Ts66Yah mice were generated as previously described [20]. Animals were housed up to 4 males and 5 females per cages (cage type : Green Line-39 x 20 x 16 cm, Techniplast, Italy) and had free access to purified water and food (D04 chow diet, Safe, Augy, *France*). The temperature in animal house was maintained at 23±1 °C, and the light cycle was controlled as 12 h light and 12 h dark (lights on at 7AM). On experimental days, animals were transferred to experimental room 30 min before the start of the test. Except for circadian activity, all experiments were performed between 8AM and 4PM. Mice were given resting time of 2 to 7 days between two different types of tests. Behavioural characterisation of a Ts66Yah mice was performed with total number of 37 animals (littermates) of both sexes (Eu= 9 males + 11 females; Ts66Yah= 8 males + 9 females) divided into 2 separate cohorts. A pipeline of behavioural tests included : circadian activity test, Open Field, Elevated plus maze, Y maze and New Object Recognition (NOR) with 1h and 24h of retention time. To avoid the effect sexual pheromones on animal behaviour, males and females were tested separately. In order to investigate an effect of ageing on cognitive behaviour, mice were tested in the same phenotyping pipeline at 3 and 9 months of age (corresponding to a start point and end point of the molecular analysis). All behavioural tests were performed in Phenotyping Platform of Mouse Clinic Institute, in accordance with the Directive of the European Parliament: 2010/63/EU, revising/replacing Directive 86/609/EEC and with French Law (Decree n° 2013-118 01 and its supporting annexes entered into legislation 01 February 2013) relative to the protection of animals used in scientific experimentation (Agreement D67-218-40 until 14/10/2028). The research project authorisation was accredited by the Ministry of National education, Higher Education and Research APAFIS#15187-201805221519333v3.

### Motor activity in circadian monitoring

Spontaneous horizontal and vertical (rears) during day/night cycle was measured in actimetric cages (Immetronic, Pessac, France). The actimeter apparatus is composed of 8 individual cages (11×21×18 cm^3^) equipped with automatic water and food distribution systems (water and food provided *ad libitum*), metal grid floor and a drawer collecting urine and feces. Each cage has 2 horizontal levels of infrared beam units connected to a computer recording beam breaks, which indicate animal’s position and horizontal and vertical movement. We used maximum of 3 sets of actimeters per experiment and never tested males and females in the same actimeter. Day/night cycle was 12h day/12h night (night cycle started at 7PM). The 45h of testing was carried out over 3 days (11AM on day1 until 8AM on day3) and was divided into 4 phases : habituation (from 11AM to 7PM on day 1), first night/dark phase (from 7PM on day 1 to 7AM on day 2), day/light phase (from 7AM to 7PM of day 2), second night/dark phase (from 7PM on day 1 to 7AM on day 3). We chose protocol with 2 night/dark phases in order to estimate animals’ circadian activity in response to new (first night/dark phase) versus familiar (second dark/night phase) environment. The test was automatically recorded using ActiTrack software. We calculated horizontal and rears activity as well as food and water intakes per hour and per test phase.

### Motor activity in the Open Field

Spontaneous locomotor activity was measured in Open Field (OF) under dim light (50lux) to diminish stress in animals. Mice were placed individually into arena (white cylindrical plastic arena; 55 cm diameter) and let to explore it for 15 min. The test was repeated 24h later to assess the adaptation/habituation to the new environment. The trials were recorded and analysed using video tracking system (Ethovision, Noldus, France). Distance travelled (m), speed, mobility and time spent in different zones of the arena were calculated. OF test was followed by Novel Object Recognition Task.

### Memory in the Novel Object Recognition

The Novel Object Recognition (NOR) task was used to evaluate recognition memory in animals previously habituated to OF arena. The NOR test was divided into 2 phases: presentation (training) and memory test. In presentation phase mice were placed individually in OF arena containing two identical objects and allowed to explore them for 10 min starting from first sniffing/touching behaviour. Both, voluntary investigation of an object at the distance less than 2cm and touching the object by an animal were considered as an exploration. Once a trial finished, mice returned to their home cage for a specific retention interval, which was 1 hour for short-term memory, and 24 hours for long-term memory examination. After the interval, mice return to the OF arena containing one familiar object (object used in presentation phase) and one novel object and are allowed to freely explore them for 10min (test phase). In this protocol, all mice were first tested for short-time memory (day1), followed by long-term memory test (training on day2 and test on day 3). Two different sets of objects were used for each of the memory tests. The percent of time exploring novel object relative to total time of exploration (novel object+ familiar object) was calculated to assess memory (RI: recognition index).). A RI of 0,5 or less indicated lack of recognition memory. Mice exploring objects less than 5 sec in either of the phase were excluded from the analysis. The trials were recorded and analysed using video tracking system (Ethovision, Noldus, France). Light intensity in experimental room during each phase of the test is 50 lux in the centre of the arena. Between each trial the arena and objects were cleaned with 50% ethanol to reduce olfactory cues. Objects were placed in intermediate zone of the arena.

### Spontaneous alternation in Y maze

Short-term working memory was evaluated by recording spontaneous alternation in the Y-maze test. The Y-maze test uses innate preference of animals that freely explore Y shaped maze (40 x 9 x16 cm, all arms at an angle of 120° to each other) to visit an arm that has not been previously explored (the wall of each arm is painted with different pattern). In a procedure, mice were placed individually in one of the arms of the maze and allowed to freely explore all three arms (named A, B, C) of the maze for 8 minutes and sequences of entries to the arms (whole body, including tail) were recorded. Between each mouse the Y maze apparatus was cleaned with 50% ethanol to reduce olfactory cues. The test was performed under bright light (100lux in the experimental room giving about 70 lux in the maze). Spontaneous alternation behaviour was calculated from successive entries of the three arms on overlapping triplet sets, in which three different arms are entered (sequence ABC, or BCA, or ACB, or BAC, or CAB, or CBA). The number of alternations was then divided by the number of alternation opportunities namely, total arm entries minus one. Deficits in spontaneous alternation occurred if frequency of alternation (ABC, etc) was lower than 50%.

### Anxiety related behaviour in the elevated plus maze

The elevated plus maze (EPM) test is commonly used for studying anxiety-like behaviour in rodents. The test is based on the natural aversion of animals for open and elevated areas, as well as on spontaneous exploratory behaviour in a new environment. The EPM apparatus (Imertonic, infrared detection system) consists of a plus-shaped maze elevated 50 cm above the floor. The maze contains two closed (wall-sheltered) and two open arms both equipped with infrared beams connected to a computer recording the beams breaks, which enable to measure movement of animals.

Test was performed under dim light (15lux). Mice were placed in the centre of the maze and let to freely explore the maze for 5 min. After each trial the EPM apparatus were cleaned with 50% ethanol to reduce olfactory cues. The preference for exploring open arms over closed arms (expressed as either a percentage of entries or a percentage of time spent in the open arms) was used to measure anxiety-like behaviour.

### Spatial learning and memory in the Morris water maze

Morris water maze experiment was performed to evaluate spatial learning and memory through a spatial search strategy that involves finding a hidden platform based on visual cues placed outside the water maze. The maze was a circular pool (150-cm diameter, 60-cm height) filled to a depth of 40 cm with water maintained at 20–22°C, opacified with white aqueous emulsion (Acusol OP 301 opacifier) and split into 4 virtual quadrants. The escape platform, made of rough plastic, was placed in the middle of one of the quadrants and submerged 1 cm below the water’s surface. Walls of the experimental room were hung with contrast drawing that served as visual cues. The room was illuminated with indirect light at 100 lux. The test consisted of two phases: learning phase and probe test. The learning phase lasted 6 consecutive days and consisted of several 90 sec trials per day, namely 3 trials per day for young 3 months old mice and 4 trials per day for 9 months old mice. Each trial started with mice facing the interior wall of the pool. The starting position was changed pseudo-randomly between trials. Trial was automatically stopped if mouse climbed onto the platform or after a maximum searching time of 90 sec. If a mouse failed to find the platform within 90 sec period, it was guided to its position by the experimenter. After each trial mice were left on the platform for 20-30 sec, and then placed under heating lamp during 10-15 min inter-trial intervals. Learning was assessed during a probe test performed 24 h after the last training session. Latency to find the platform, distance travelled in each quadrant and the average speed were quantified. During the probe test the target platform was removed from the pool and mice were allowed to swim in the maze for 60 sec. A percentage of time spent in platform quadrant demonstrated the memory with 25 % or less indicating no memory. To distinguish an impairment in spatial learning versus ocular defects, few days after the Morris water maze task, we performed a cued version test that consisted of 4 trials of 60s, in which the platform was labelled by a flag and its location varied between the trials. The test was recorded and analysed using video tracking system (Ethovision, Noldus, France).

### Protein Sample Preparation

The total protein extract from Ts66Yah and Eu mice at 3,6 and 9 months of age (12 mice per age group, 6 females and 6 males) was prepared in RIPA buffer (pH 7.4) containing tris-HCl (50 mM pH = 7.4), NaCl (150 mM), 1% NP-40, 0.25% sodium deoxycholate, 1 mM EDTA, 0.1% SDS, supplemented with phosphatase and protease inhibitor (539132, Millipore, 1:100; P0044; Sigma-Aldrich, St. Louis, MO, USA; 1:100). Before clarification, the samples were sonicated on ice and then centrifuged at 14,000× rpm at 4 ◦C for 30 min to remove cellular debris. The supernatant was then used to determine the total protein concentration by the BCA method (Pierce, Rockford, IL, USA).

### Western blot

For western blots, 15 µg of proteins were resolved on Criterion TGX Stain-Free 4-15%,18 and 26-well gel (Bio-Rad Laboratories, #5678084, #5678085) in a Criterion large format electrophoresis cell (Bio-Rad Laboratories, #1656001) in Tris/Glycine/SDS (TGS) Running Buffer (Bio-Rad Laboratories, #1610772). Immediately after electrophoresis, the gel was transferred on a Chemi/UV/Stain-Free tray and then placed into a ChemiDoc MP imaging System (Bio-Rad Laboratories, #17001402) and UV-activated based with Image Lab Software (Bio-Rad Laboratories) to collect total protein load image. Following electrophoresis and gel imaging, the proteins were transferred via the TransBlot Turbo semi-dry blotting apparatus (Bio-Rad Laboratories, #1704150) onto nitrocellulose membranes (Bio-Rad, Hercules, CA, USA, #162-0115). The membranes were blocked with 3% bovine serum albumin in 0.5% Tween-20/Tris-buffered saline (TTBS) and incubated overnight at 4 °C with the antibodies reported in supplementary table 1. After 3 washes with Tween-20/Tris buffered saline (TTBS) buffer the membranes were incubated for 1 hour at room temperature with anti-rabbit/mouse/goat IgG secondary antibody conjugated with horseradish peroxidase (1:5000; Sigma–Aldrich, St Louis, MO, USA). Signals was detected with Clarity enhanced chemiluminescence (ECL) substrate (Bio-Rad Laboratories, #1705061), then acquired with Chemi-Doc MP (Bio-Rad, Hercules, CA, USA) and analysed using Image Lab software (Bio-Rad, Hercules, CA, USA).

### Slot blot

For total Protein Carbonyls (PC) levels: hippocampal total protein extract samples (5 µl), 12% sodium dodecyl sulphate (SDS; 5 µl), and 10 µl of 10 times diluted 2,4-dinitrophenylhydrazine (DNPH) from 200 mM stock were incubated at room temperature for 20 min, followed by neutralization with 7.5 µl neutralization solution (2 M Tris in 30% glycerol) and then loaded onto nitrocellulose membrane as described below. For total (i) protein-bound 4-hydroxy-2-nonenals (HNE) and (ii) 3-nitrotyrosine (3-NT) levels: hippocampal total protein extract samples (5 µl), 12% SDS (5 µl), and 5 µl modified Laemmli buffer containing 0.125 M Tris base, pH 6.8, 4% (v/v) SDS, and 20% (v/v) glycerol were incubated for 20 min at room temperature and then loaded onto nitrocellulose membrane as described below. Proteins (250 ng) were loaded in each well on a nitrocellulose membrane under vacuum using a slot blot apparatus. The membranes were blocked with 3% bovine serum albumin in 0.5% Tween-20/Tris-buffered saline (TTBS) for 1 h and incubated with an anti-2,4-dinitrophenylhydrazone (DNP) adducts polyclonal antibody (1:100, EMD Millipore, Billerica, MA, USA, #MAB2223) or HNE polyclonal antibody (1:2000, Novus Biologicals, Abingdon, United Kingdom, #NB100-63093) or an anti 3-NT polyclonal antibody (1:1000, Sigma-Aldrich, St Louis, MO, USA, # N5538) for 90 min. The membranes were washed in PBS following primary antibody incubation three times at intervals of 5 min each. The membranes were incubated after washing with an anti-rabbit IgG alkaline phosphatase secondary antibody (1:5000, Sigma– Aldrich, St Louis, MO, USA) for 1 h. The membranes were washed three times 5 min in PBS and developed with Sigma fast tablets (5-bromo-4-chloro-3-indolyl phosphate/nitroblue tetrazolium substrate [BCIP/NBT substrate]). Blots were dried, acquired with Chemi-Doc MP (Bio-Rad, Hercules, CA, USA) and analysed using Image Lab software (Bio-Rad, Hercules, CA, USA). No non-specific binding of antibody to the membrane was observed.

### Subcellular fractionation

Subcellular fractionation was prepared using the hippocampus of Ts66Yah an Eu mice (3,6 and 9 months, 6 mice per group). To perform the isolation protocol, we used 10 mg of hippocampal tissue. We used three female and three male belonging to each experimental group. The hippocampal tissue was transferred into Eppendorf tubes containing STM buffer supplemented by the addition of protease inhibitor cocktail (1:100; 539132, Millipore) and phosphatase inhibitor cocktail (1:100; P5726, Sigma-Aldrich, St Louis, MO, USA) and homogenized on ice using a tight-fitting Teflon pestle. All the steps for subcellular fractions separation are in accordance with those reported by Dimauro et al., (2012) [22]. Validation of the purity of the subcellular fractions was determined by examining DNA Polymerase II in the nuclear fraction and Complex I in the mitochondrial fraction by Western Blot analysis.

### Statistical Analysis

Behavioural data are presented as mean group value± standard error of the mean (SEM). Normal distribution was evaluated with Shapiro-Wilk test. Differences were considered significant at p<0.05. As no difference between males and females were present, data were pooled. We performed unpaired two-tailed t-test between genotypes or repeated measures Anova (genotype, age) with Bonferroni post hoc test. One sample t-test for: (i) NOR test to evaluate recognition index against chance level (50%), (ii) in MWM task to compare performance against chance level (25%), (iii) Y maze to evaluate spontaneous alternation behaviour against chance level 50%. For the multivariate analysis we identified the differentiating variables allowing to separate the young and old mouse from the two genotypes by using the GDAPHEN pipeline [23]. Briefly, we used 12 selected variables, plus the sex and genotype, derived from the mouse behavioural dataset with the groups of same individuals analysed at 3 and 9 months of age.

Molecular Data were calculated as a percentage of Eu at 3 month and are expressed as mean ± SEM. We performed both western blots and slot blots analyses by comparing Eu and Ts66Yah mice at each age (3,6 and 9), separately. With the aim to compare all the groups of analysis, we performed western blot analysis including only Eu mice of all the age of analysis. Hence, data were normalized for the average of Eu mice. Then, data were correlated all together and normalized for the average of Eu mice at 3 months. 2-way analyses of variance (Anova) with diagnostic group (Eu vs Ts66Yah) followed by Fisher LSD test was used. Data are presented as means ± SEM. Columns were used to show differences among the groups (Eu vs Ts66Yah). Dots were used for Eu and triangle for Ts66Yah. White colour indicates female mice (n =6) and black colour indicate male mice (n =6). Lines were used to show age-associated changes within each group. For columns: *p < 0.05, **p < 0.01, ***p < 0.001 and ****p < 0.0001 (2-way Anova with Fisher’s LSD test). For lines: #p < 0.05, ##p < 0.01, ###p < 0.001 and ####p < 0.0001 among 3 months vs 6 and 9 months; +p < 0.05, ++p < 0.01, +++p < 0.001 and ++++p < 0.0001 among 6 months vs 9 months (2-way Anova with Fisher’s LSD test). To evaluate the effects of genotype, aging and sex, and of their interactions, on collected data we performed a 3-way ANOVA analysis. Statistical analyses and graphs for both behavioural and molecular data were done using GraphPad Prism 10 software for univariate analysis.

## RESULTS

### Aging effect on behavioural performance of the Ts66Yah DS mouse model

To investigate the effect of aging on the Ts66Yah DS mouse model, we produced a cohort of DS mouse models with their control littermates, including males and females, in a balanced ratio, that were studied at young age (3 months) and latter at 9 months of age (adult). We did not notice any welfare issue during this aging protocol. This cohort was investigated for circadian activity, open field exploration, object recognition, alternation, exploration of the elevated plus maze, and the Morris water maze to study locomotor activity, object memory, anxiety related behaviour and spatial memory. All these tests were done over 6 weeks at each time point. We found no significant alteration in the motor activity (horizontal and vertical) during circadian monitoring and open field exploration (Figure S1). Both short-term (1h) and long-term (24h) object discrimination memory (scored with the Recognition Index; RI) was defective in the Ts66Yah mouse model at the age of 3 months (RI=0,51 for short-term and RI=0,49 for long-term recognition memory), comparing to age-matched Eu control littermates (RI=0,66 for short term and RI=0,57 for long-term recognition memory). At the age of 9 months both short-term and long term-recognition memory was defective in Ts66Yah mice (RI=0,52 and RI=0,48, respectively) (Figure 1A-B). We observed a reduced object memory with a 24h retention time in the Eu littermate group at 9 months of age (RI=0,51), which could probably be explained by a batch effect, since the short-term memory is preserved in these mice (RI=0,65). The working memory was impaired at both ages in the Ts66Yah mice (alternation of 54% and 50 % for 3- and 9-months old animals, respectively), whereas it was not affected in control littermates (alternation of 62% and 59% for 3 months old and 9 months old Eu, respectively; Figure 1C). These results agreed with our original publication of the Ts66Yah DS mouse model, but extended the defects to adult during ageing [20].

**Figure 1.**
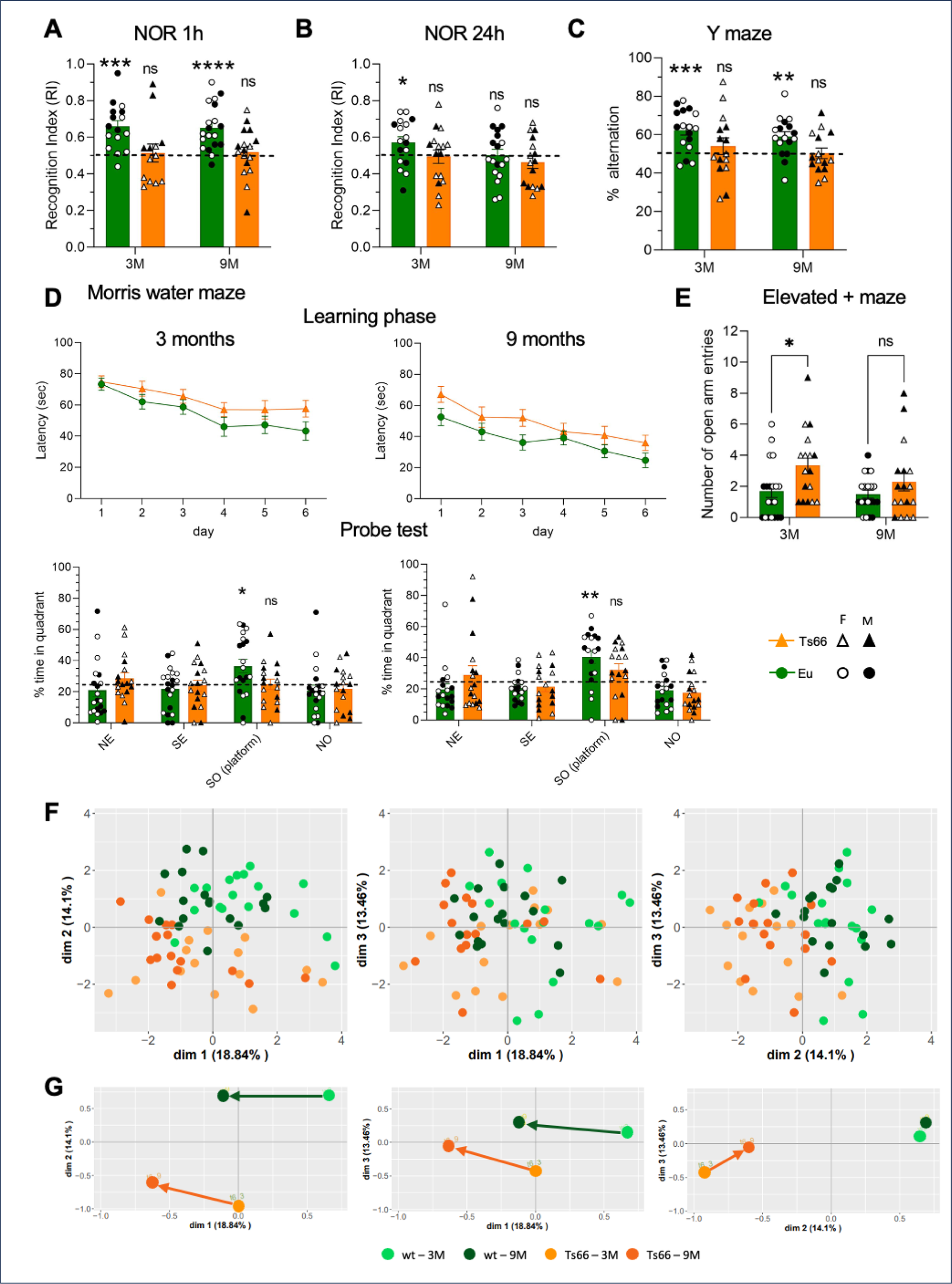
Cognitive and behaviour deficits in Ts66Yah mice during ageing. Short-term recognition memory in Ts66Yah mice in NOR test with 1h retention time **(A)** and 24h retention time **(B)** at 3 months and 9 months of age (RI=new T /(new + familiar) T; statistics: one sample t test, chance level 0,5). Ts66Yah mice display working memory deficits in Y maze test **(C)** (statistics: one sample t test, chance level 50%). In MWM, there was no significant difference between Ts66Yah and Eu mice in a latency to reach platform in learning phase **(D)**, although in a probe test Ts66Yah mice showed memory deficits at both, 3 months and 9 months of age (statistics: one sample t test, chance level 25%).; 3 months old Ts66Yah mice display anxiolytic-like behaviour in EPM test as measured by the number of the open arms visits **(E)** (* p=0,05). All data are presented as means ± SEM. **(F)** 2D projection of the multifactorial analysis of the Ts66Yah mouse compared to control littermates (Eu) with 11 non-correlated behavioural variables assessed at 3- and 9-months unravelled evolution of trajectory along ageing with similar or specific directions for each genotype from 3 to 9 months **(G)**.

To go further, we decided to modify the learning phase of the Morris water maze for young 3 months old mice, making the learning more difficult/challenging with only 3 sessions instead of 4 as used previously in Duchon et al. [20]. We were keen on observing, with this reduced training, a spatial memory deficit in the probe test (measured by % of time spent in a platform quadrant) in 3 months-old Ts66Yah (24,9%) compared to Eu littermates (36,4%) (Figure 1D). A similar memory deficit was found in the 9 months-old DS mice (32,1 % in Ts66Yah comparing to 40,5% in Eu). Thus, with a more challenging Morris water maze protocol, we found an impairment of the spatial memory in the Ts66Yah mouse models as early as in 3 months old mice. We also explored the anxiety related behaviour with the elevated plus maze (Figure 1E). Increased number of open arm entries was detected in the Ts66Yah mouse individuals compared to Eu littermate at 3 months of age (3,4 in Ts66Yah to 1,7 in Eu, p=0,02), but the phenotype was not found significative at 9 months (2,3 in Ts66Yah to 1,5 in Eu). The behavioural analysis did not show any significant sex difference for the tests performed, therefore the data was pooled.

To discriminate the genotype and sex effects in this behavioural study, we performed a multivariate analysis using the GDAPHEN pipeline with the same 12 variables assessed along ageing [23]. As observed in Figure 1F, the two genotypes were easily separated in the first 3 dimensions, considering 46,5% of the total variance (Supplementary Figure 2A-B). Interestingly different non-correlated variables were contributing to defining the first 3 dimensions (Supplementary Figure 2C). Three variables derived from the elevated plus maze test, brought the main contributions to dimension 1 (Supplementary Figure 2D). For dimension 2, the NOR variables were the main differentiating variables aligned with the genotype while for dimension 3 two variables from the Morris water maze aligned with the sexes. Looking more precisely at the centroid of each group of points in each 2D representation (Figure 1G), we observed that the directionality from 3 to 9 months is preserved between the two genotypes in dimensions 1 and 2, suggesting a common path driven by ageing on these variables affecting both genotypes. Nevertheless, dimension 3 showed a specific difference in the trajectory of the centroid from 3 to 9 months in the Ts66Yah. The two variables “Distance to find the platform” and the “velocity” in probe test were affecting the trajectory of both sexes during ageing with almost no effect in Eu. This last observation suggested that a specific effect on ageing is found in the Ts66Yah genotype compared to Euploid control.

Overall, the behavioural features of the Ts66Yah mice were confirmed in this new assessment with a major improvement to detect spatial memory defect and also new anxiety related phenotype in young individuals. Interestingly the mean ageing trajectory based on multivariate analysis unravelled two main directionalities shared between the Ts66Yah individuals and the Euploid control and one small but specific ageing effect only affecting both sexes with the Ts66Yah genotype.

### Molecular signatures of neurodegeneration are evident in Ts66Yah mice

Individuals with DS have a higher risk to develop AD during aging. To unravel the molecular mechanisms underlying brain degeneration in Ts66Yah mice, we initially examined the neuropathological signatures of AD, known to develop in DS compared to Eu mice [24]. We took advantages of the cohort from the behavioural analysis, to isolate samples for the biochemical analysis, and we also included a new group of mice at 6 months of age to observe the molecular transition from young to adult mice. We investigated the molecular signature using western blot analysis for a selection of targets. To elucidate both genotype- and age-related effects, we present the molecular findings through two distinct graphs (both derived from the same dataset). In the first graph (columns), we emphasize genotype differences between 3, 6, and 9 months (Eu vs. Ts66Yah), while in the second graph (dots), we highlight aging-related changes within each genotype group. We initially assessed the levels of the APP, and our data confirmed the expected up-regulation in mice at 3 months, 6 months, and 9 months of age; a direct consequence of the copy number increase of the *App* gene in the Ts66Yah mice (Figure 2 B and B1). Interestingly the APP level at 3 months in the Ts66Yah mice was almost similar to wild-type mice at 6 and 9 months (Figure 2B1). A significant influence of both genotype [F (1, 60) = 162.9, P<0.0001] and age [F (2, 60) = 43.93, P<0.0001] is demonstrated for APP. The phosphorylation of APP at T668 was shown to increase Aβ levels by facilitating exposure to BACE cleavage [25]. We found a significant increase of T668 phosphorylation in Ts66Yah mice starting at 6 months of age and persisting in 9 months-old mice compared to 3 months (Figure 2 C and C1). The overexpression and hyperphosphorylation of TAU, specifically on serine and threonine residues, leads to protein self-assembly and the formation of toxic neurofibrillary tangles (NFTs), a well-established hallmark of AD-like pathology [26]. Data from Ts66Yah mice indicated a significant increase in total TAU at 3, 6 and 9 months old when compared to age-matched Eu mice (Figure 2 E and E1). As observed for APP, the level of TAU in Ts66Yah brain at 3 months is similar to 9 months-old control animals. A significant influence of genotype [F (1, 60) = 31.93, P<0.0001] and aging [F (2, 60) = 10.02, P=0.0002] was detected for total TAU levels. Phosphorylation of TAU on both Ser202-Thr205 residues (AT8) in Ts66Yah mice demonstrated a significant increase at 6 at 9 months (Figure 2F). Changes in Ser202-Thr205 within the groups showed a significant variability with age in Ts66Yah, particularly with an increase from 3 to 6 months, and a decrease from 6 to 9 (Figure 2 F1). We further examined DYRK1A, a kinase involved in TAU phosphorylation in AD pathogenesis, whose gene is triplicated in the DS context. DYRK1A protein levels were elevated in Ts66Yah mice compared to Eu mice at 3, 6 and 9 months of age (Figure 2D). Moreover, age-related changes in DYRK1A within each group highlighted that Ts66Yah mice exhibited a significant increase from 3 to 9 months old (Figure 2 D1). DYRK1A up-regulation is associated with both age [F (2, 59) = 14.40, P<0.0001] and genotype [F (1, 59) = 85.72, P<0.0001]. Similar to APP and TAU, DYRK1A levels increases in Ts66Yah mice from 3 to 9-months of age. Altogether the well-known landmarks of AD were found overexpressed in the Ts66Yah brain already at 3 months and most of them continued to accumulate during aging.

**Figure 2.**
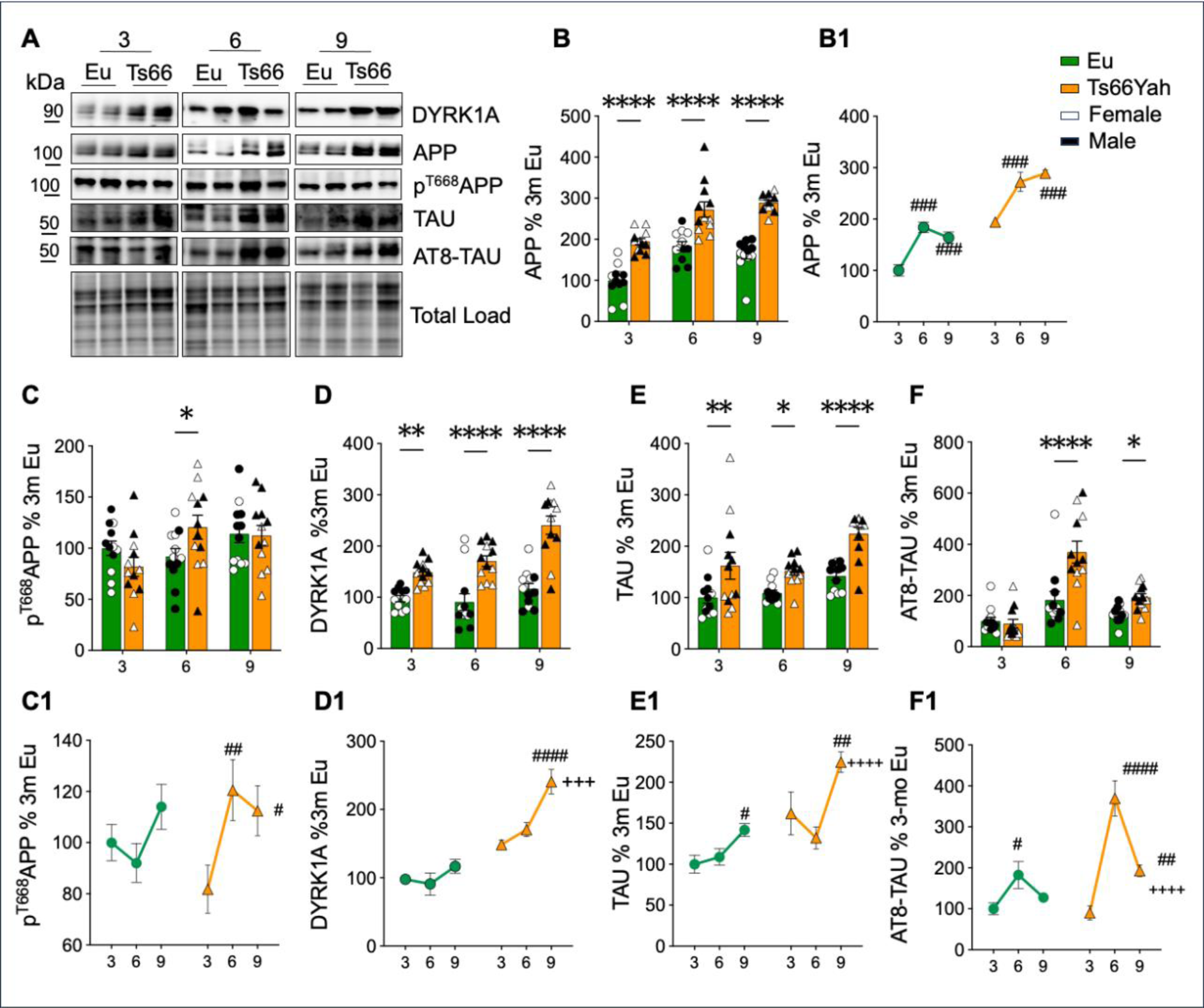
Evaluation of age- and genotype-associated effects in major molecular hallmarks of neurodegeneration in Ts66Yah mice. **(A)** Representative Western blot images and densitometric evaluation of APP full length **(B and B1)**, p ^T668^APP (**C and C.1**), DYRK1A **(D and D1),** TAU (**E and E1**) and phosphorylated TAU on residues S202 and T205 (AT8) (**F and F1**) protein levels were evaluated in the hippocampus of Eu and Ts66Yah mice at 3 months (Eu n = 12, Ts66Yah n = 12); 6 months (Eu n = 12, Ts66Yah n = 12); 9 months (Eu n = 12, Ts66Yah n = 12). Protein levels were normalized per total protein load. All densitometric values are given as percentage of Eu at 3 months set as 100%. Data are presented as means ± SEM. Columns were used to show differences among the groups (Eu vs Ts66Yah). Dots were used for Eu and triangle for Ts66Yah. White colour indicates female mice (n =6) and black colour indicate male mice (n =6). Lines were used to show age-associated changes within each group. For columns: *p < 0.05, **p < 0.01, ***p < 0.001 and ****p < 0.0001. For lines: #p < 0.05, ##p < 0.01, ###p < 0.001 and ####p < 0.0001 among 3 months vs 6 and 9 months; +p < 0.05, ++p < 0.01, +++p < 0.001 and ++++p < 0.0001 among 6 months vs 9 months (2-way ANOVA with Fisher’s LSD test).

### The alteration of antioxidant responses promotes unbalanced redox homeostasis in Ts66Yah mice

Redox unbalance is an early event in subject with DS and in transgenic mice of the disease [27]. An essential antioxidant gene, whose triplication is involved in the redox imbalance evident in the DS context, is *SOD1* localised in 21q22.1. SOD1 is responsible for catalysing dismutation of superoxide to hydrogen peroxide and molecular oxygen. As expected from a gene dosage effect, the levels of SOD1 were up regulated in Ts66Yah mice hippocampus with respect to Eu mice at 3, 6 and 9 months (Figure 3E and E1). A strong effect for genotype [F (1, 60) = 43.31, P<0.0001] and sex [F (1, 60) = 12.39, P=0.0008] was observed for SOD1. To further elucidate the dysregulation of antioxidant responses in Ts66Yah mice, we investigated the NRF2 signal. To note BACH1, triplicated in DS, engages a competition with NRF2 for the regulation of the ARE-mediated antioxidant enzymes [28]. The analysis performed on the nuclear subcellular fraction highlighted a significant decrease of NRF2 in 3-month-old and 9-month-old Ts66Yah mice (Figure 3F and F1). On the contrary, BACH1 protein was significantly up regulated in 6 months old Ts66Yah (Figure 3G and G1). The analysis of the nuclear BACH1/NRF2 ratio may serve as an indicator of antioxidant response efficiency. The observed increase of BACH1/NRF2 at 6 and 9 months of age in Ts66Yah mice support a potential inhibition of the pathway (Figure 3H and H1). BACH1/NRF2 changes are associated with age [F (2, 24) = 43.85, P<0.0001] and genotype [F (1, 24) = 91.84, P<0.0001]. To further confirm the reduced induction of the NRF2-related antioxidant response, we analysed HO-1 expression, and we observed a significant decrease at 3 months and 9 months of age in Ts66Yah mice (Figure 3I and I1). HO-1 changes are influenced by genotype [F (1, 60) = 6.407, P=0.0140]. The analysis of markers of protein oxidative damage, namely protein nitration (3-NT), 4-hydroxy-2-nonenal (HNE) protein adducts and protein carbonyls (PC) were performed in Ts66Yah mice to account for the increase of oxidative stress expected in DS phenotype. DS animals demonstrated a consistent increase in 3-NT at 6 months and 9 months, but no significant changes in 3-month-old mice was observed (Figure 3A). Conversely, immunochemical analysis of HNE showed significant alterations at 3, 6 and 9 months of age (Figure 3B). Consistently, the analysis of carbonylated proteins showed a significant increased accumulation of PC at 6 and 9 months of age (Figure 3C and C1). Grouped analysis revealed a more pronounced elevation of both 3-NT and PC in Ts66Yah mice as they aged, compared to Eu mice (Figure 3A1, 3B1 and 3C1) supporting for both the influence of genotype and aging (See Figure 10A).

**Figure 3.**
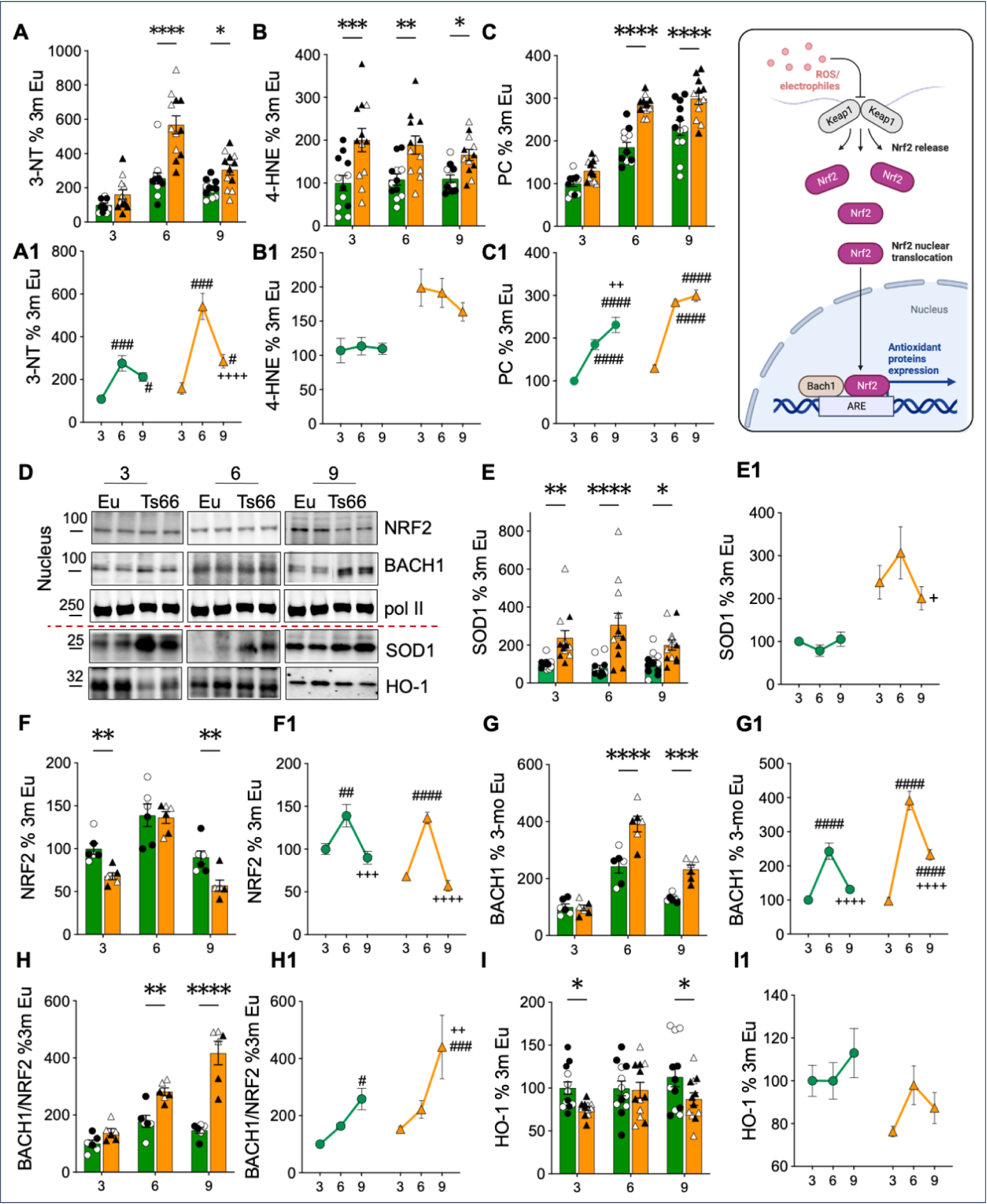
Investigation of oxidatively damaged proteins and antioxidant responses in the hippocampus of Ts66Yah. Slot Blot quantification of 3-NT levels (**A and A1**), 4-hydroxy-2-nonenal (HNE) **(B and B1)** and protein Carbonyls **(PC; C and C1)** in the hippocampus of Eu and Ts66Yah mice at different ages 3 months (Eu n = 12, Ts66Yah n = 12); 6 months (Eu n = 12, Ts66Yah n = 12); 9 months (Eu n = 12, Ts66Yah n = 12). Representative Western blot images and densitometric evaluation of SOD1 (**E and E1**) and HO-1 (**I and I1**) protein levels in Eu and Ts66Yah mice at 3 months (Eu n = 12, Ts66Yah n = 12), 6 months (Eu n = 12, Ts66Yah n = 12) and 9 months (Eu n = 12, Ts66Yah n = 12). Western blot analysis and densitometric evaluation showing protein levels of nuclear NRF2 (**F and F1**), BACH1 (**G and G1**) and BACH1/NRF2 ratio (**H and H1**) in the hippocampus of Eu and Ts66Yah mice at 3 months (Eu n = 6, Ts66Yah n = 6); 6 months (Eu n = 6, Ts66Yah n = 6); 9 months (Eu n = 6, Ts66Yah n = 6). Protein levels were normalized per total protein load. All densitometric values are given as percentage of Eu at 3 months set as 100%. Data are presented as means ± SEM. Columns were used to show differences among the groups (Eu vs Ts66Yah). Dots were used for Eu and triangle for Ts66Yah. White colour indicates female mice and black colour indicate male mice. Lines were used to show age-associated changes within each group. For columns: *p < 0.05, **p < 0.01, ***p < 0.001 and ****p < 0.0001. For lines: #p < 0.05, ##p < 0.01, ###p < 0.001 and ####p < 0.0001 among 3 months vs 6 and 9 months; +p < 0.05, ++p < 0.01, +++p < 0.001 and ++++p < 0.0001 among 6 months vs 9 months. In (C), scheme relative to the portion of the pathway mentioned in this figure. Created with Biorender.com.

Remarkably, the Ts66Yah mice displayed constitutive overexpression of SOD1, as anticipated due to dosage effect with increased markers of protein oxidative damage, while the brain overexpression of BACH1 was only detected at 6 months of age, when NRF2 was also found overexpressed. Oppositely, NRF2 levels were decreased in 3- and 9-month-old Ts66Yah brain with a concomitant reduction of HO-1 expression. Here too, the contribution of DS related gene overdosage was correlated with unbalanced redox activities in Ts66Yah.

### Brain metabolism: The alteration of components of insulin signalling and mitochondrial OXPHOS support metabolic failure and redox unbalance in Ts66Yah mice

Deficiencies in redox homeostasis are associated with impaired brain energy metabolism due to the increased ROS production leaked by defective mitochondria. Therefore, to investigate mitochondrial and OXPHOS status we evaluated changes for the levels of electron transport chain (ETC) Complexes (I-IV) and Complex V (ATP synthase) in mitochondrial extracts from Ts66Yah and Eu mice at different ages. We observed that at 3 months of age Ts66Yah mice exhibited a consistent reduction for Complex I level (Figure 4B). This reduction in complex I was also evident as both Eu and Ts66Yah mice aged. At 6 months of age, there was a trend towards reduced levels in all complexes, with a significant reduction in Complex III (Figure 4D). However, the most pronounced decrease in mitochondrial protein levels was observed in adult Ts66Yah mice. Indeed, 9 months old Ts66Yah demonstrated a significant reduction in Complex I, Complex II, Complex IV and Complex V levels (Figure 4B-C-E and F). These findings suggest a close connection between mitochondrial defects and oxidative stress levels in the brains of Ts66Yah mice. Collectively results on mitochondrial complexes demonstrate an altered expression associated with both genotype and aging but not with sex (see 3-way ANOVA data in Figure 10A).

**Figure 4.**
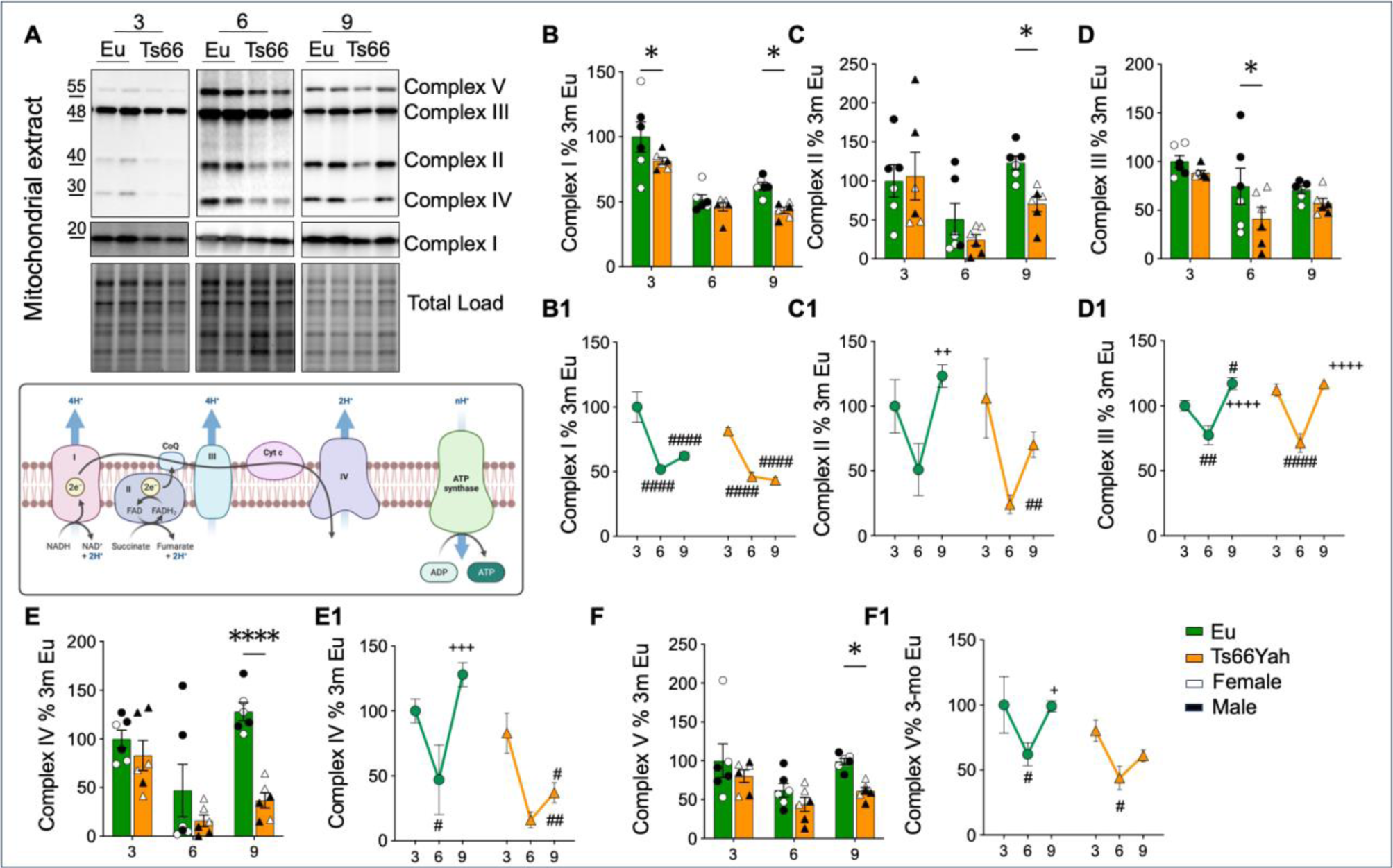
Evaluation of protein components of the mitochondrial ETC machinery in Ts66Yah hippocampus. **(A)** Representative Western blot images and densitometric evaluation of Complex I **(B and B1)**, Complex II **(C and C1)**, Complex III **(D and D1)**, Complex IV **(E and E1)** and complex V **(F and F1)** in the mitochondrial extract of Eu and Ts66Yah mice hippocampus at different ages 3 months (Eu n = 6, Ts66Yah n = 6); 6 months (Eu n = 6, Ts66Yah n = 6); 9 months (Eu n = 6, Ts66Yah n = 6). Protein levels were normalized per total protein load. All densitometric values are given as percentage of Eu at 3 months set as 100%. Data are presented as means ± SEM. Columns were used to show differences among the groups (Eu vs Ts66Yah). Dots were used for Eu and triangle for Ts66Yah. White colour indicates female mice (n =6) and black colour indicate male mice (n =6). Lines were used to show age-associated changes within each group. For columns: *p < 0.05, **p < 0.01, ***p < 0.001 and ****p < 0.0001. For lines: #p < 0.05, ##p < 0.01, ###p < 0.001 and ####p < 0.0001 among 3 months vs 6 and 9 months; +p < 0.05, ++p < 0.01, +++p < 0.001 and ++++p < 0.0001 among 6 months vs 9 months. In (B1), scheme relative to the portion of the pathway mentioned in this figure. Created with Biorender.com.

To investigate insulin signalling we focus the analysis on the activatory state of the insulin receptor (IR) and of the insulin receptor substrate 1 (IRS1). The activation of IR, represented as p^Y1158,1162,1163^/IR, was notably elevated at 9 months in Ts66Yah mice compared to their Eu counterparts (Figure 5B and B1). When examining changes within each group as they aged, it became evident that in Ts66Yah mice, IR activation significantly increased in comparison to Eu mice. This increase was more pronounced in Ts66Yah mice than in Eu mice. Significant variations were observed particularly from 3 to 9 months and 6 to 9 months (Figure 4B1; 3-way Anova analysis showed a strong age influence [F (2, 60) = 71.98, P<0.0001]. Data sets were also influenced by genotype [F (1, 60) = 6.183, P=0.0157] and sex [F (1, 60) = 10.62, P=0.0018]). Moving downstream from IR, we found that IRS1 protein levels did not exhibit significant differences between Ts66Yah and Eu mice (Supplementary Figure 3B). Instead, it displayed age-related changes within each respective group (Supplementary Figure 3B1). Interestingly we observed a significant increase in the inhibition of IRS1, evaluated by the ratio S636/Y632, in Ts66Yah mice at 6 and 9 months old Ts66Yah (Figure 5C and C1). Changes of S636/Y632 are associated with age [F (2, 60) = 17.45, P<0.0001] and genotype [F (1, 60) = 9.784, P=0.0027]. We then evaluated whether the inhibition of IRS1 in the hippocampus of Ts66Yah mice was associated with defects in the PI3K-AKT axis. Specifically, while AKT protein levels did not change in Ts66Yah compared to Eu mice (Figure S3F), AKT activation (p^S473^/AKT) was higher in the brains of Ts66Yah compared to that in Eu mice at 9 months of age (Figure 5E). Changes in AKT activation within the groups showed a significant and more pronounced instabilities with age in Ts66Yah, particularly from 3 to 6 months, 3 to 9 and 6 to 9 months (Figure. 5E1; 3-way Anova analysis demonstrates a significant influence of age [F (2, 60) = 24.44, P<0.0001]). AKT activation was independent from PI3K induction that did not show any significant differences between Eu and Ts66Yah (Figure 5D-D1 and S3E-E1). As GSK3β is a direct target of AKT, we assessed the phosphorylation status of S9, a crucial site for inhibiting GSK3β. As illustrated in Figure 5F, the S9 phosphorylation of GSK3β showed no significant genotype differences across all age groups considered. Changes in GSK3β activation within the groups showed significant fluctuations with age in Eu and Ts66Yah (Figure 5 F1; 3-way Anova analysis [F (2, 60) = 155.4, P<0.0001]). In conclusion several mitochondrial protein complexes were down-regulated in aged Ts66Yah (9 months), while insulin signalling data suggest an uncoupling between IR, IRS1 and downstream events.

**Figure 5.**
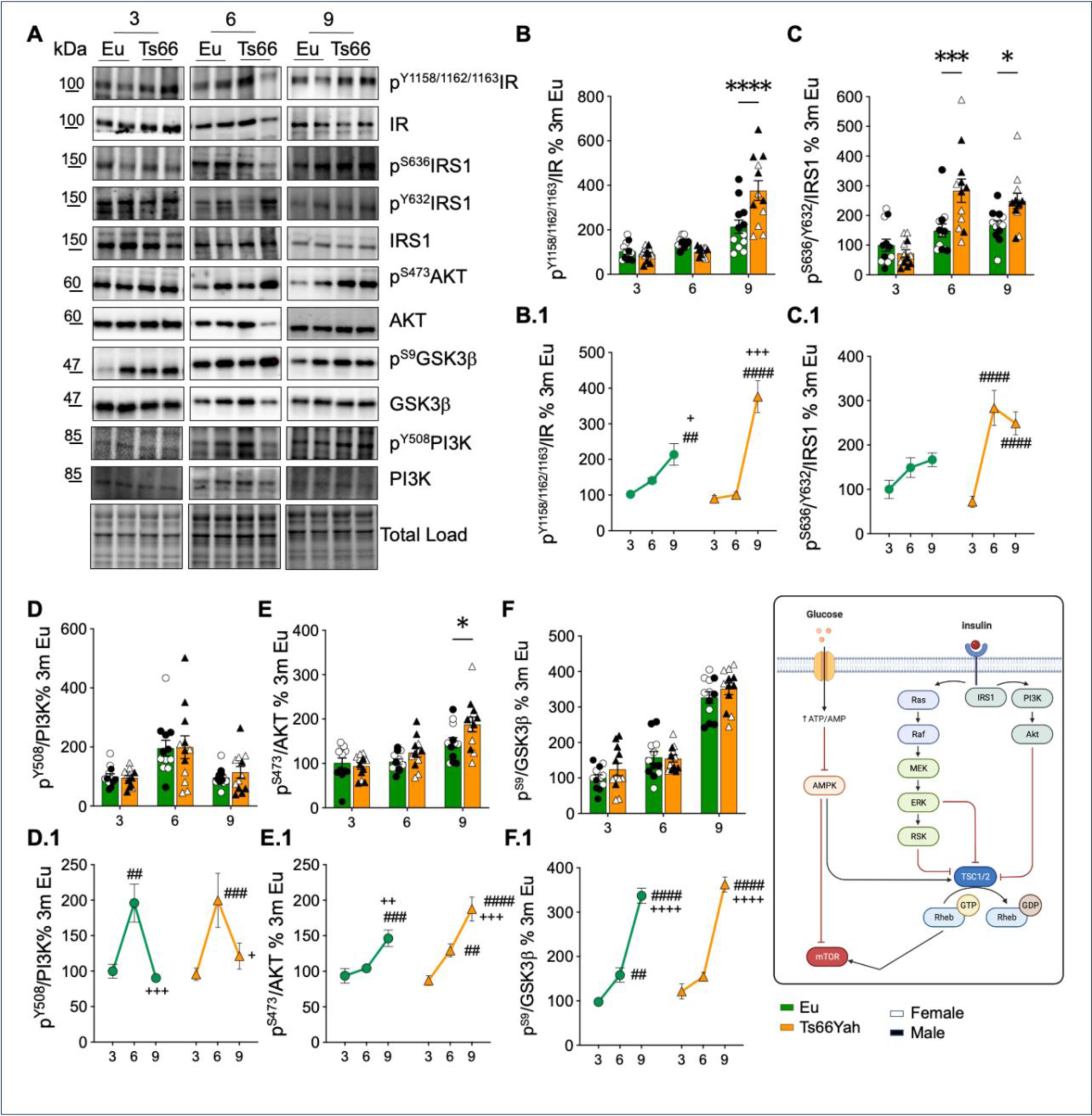
Investigation of Insulin cascade in Ts66Yah hippocampus. Representative western blot images **(A)** and densitometric evaluation of (**B and B1**) IR activation (p^Y1158/1162/1163^/IR), (**C and C1**) IRS1 inhibition (p^S636/Y632^/IRS1), (**D and D1**) PI3K activation (p^Y508^/PI3K), (**E and E1**) AKT activation (p^S473^/AKT) and (**F and F1**) GSK3β inhibition (p^S9^/GSK3β) in the hippocampus of Eu and Ts66Yah mice at different ages 3 months (Eu n = 12, Ts66Yah n = 12); 6 months (Eu n = 12, Ts66Yah n = 12); 9 months (Eu n = 12, Ts66Yah n = 12). Protein levels were normalized per total protein load. IR-, IRS1-, PI3K-, AKT- and GSK3β-associated phosphorylation were normalized by taking into account the respective protein levels and are expressed as the ratio between the phosphorylated form and the total protein levels: p^Y1158/1162/1163^/IR; p^S636/Y632^/IRS1; p^S473^/AKT; p^Y508^/PI3K; and p^S9^/GSK3β. All densitometric values are given as percentage of Eu at 3 months set as 100%. Data are presented as means ± SEM. Columns were used to show differences among the groups (Eu vs Ts66Yah). Dots were used for Eu and triangle for Ts66Yah. White colour indicates female mice (n =6) and black colour indicate male mice (n =6). Lines were used to show age-associated changes within each group. For columns: *p < 0.05, **p < 0.01, ***p < 0.001 and ****p < 0.0001. For lines: #p < 0.05, ##p < 0.01, ###p < 0.001 and ####p < 0.0001 among 3 months vs 6 and 9 months; +p < 0.05, ++p < 0.01, +++p < 0.001 and ++++p < 0.0001 among 6 months vs 9 months. In (D), scheme relative to the portion of the pathway mentioned in this figure. Created with Biorender.com.

### The impairment of proteostasis network is demonstrated by reduced autophagy and by the aberrant induction of the unfolded protein response

The loss of proteostasis is an early event in DS human brain and it correlates with increased oxidative damage [28]. To investigate if these alterations are conserved in the Ts66Yah mice, we evaluated the mTOR/autophagy axis and the induction of the PERK pathway of the unfolded protein response (UPR). mTOR is a master regulator of autophagy and sustained mTORC1 activation, indicated by S2448 phosphorylation, is known to mediate a decline in autophagy function by interacting with components involved in autophagosome formation [29]. At 3 months of age, we found no significant differences in mTORC1 activation, while a significant increase was observed at 6 and 9 months in Ts66Yah mice (Figure 6B). Notably, the changes in mTORC1 activation exhibited a distinct profile in Ts66Yah compared to Eu mice, with a significant rise observed only in Ts66Yah. Particularly the increase was observed from 3 to 6 months (Figure 6B and B1). Age, sex and genotype significantly influence the alterations in mTORC1 activation (age: [F (2, 60) = 12.06, P<0.0001]; sex: [F (1, 60) = 4.102, P=0.0473]; genotype: [F (1, 60) = 7.922, P=0.0066)]). The AMP-activated protein kinase (AMPK) serves as a key energy sensor, regulating cellular metabolism to maintain energy balance. AMPK acts upstream of mTOR, and its activation inhibits mTORC1 signalling, suppressing protein synthesis to conserve cellular energy during low energy states. We observed increased expression of AMPK at 6 and 9 months in Ts66Yah mice (Supplementary Figure 4B and B1) but a reduced AMPK activation (indicated by the p^T172^/AMPK ratio) at 9 months in Ts66Yah mice compared to Eu mice (Figure 6C). Changes in AMPK activation displayed a different pattern in Ts66Yah mice compared to Eu mice, with a significant drop observed during aging in Ts66Yah mice. A significant decrease was observed between 6 and 9 months in Ts66Yah mice (Figure 6C1). The reduction in AMPK activation was primarily driven by a significant increase in AMPK protein levels (Supplementary Figure 4B). Both mTORC1 and AMPK regulates, upon activation, the induction of autophagy through the phosphorylation of ULK1. Phosphorylation of ULK1 at S317 and S777 by AMPK induce autophagy, while phosphorylation of ULK1 at S757 by mTORC1 abolish its interaction with AMPK and reduce autophagosome formation. We observed a significant increase in ULK1 inhibition (p^S757^/ULK1) at 3 months of age in Ts66Yah (Figure 6D). However, at 6 and 9 months, we did not observe any statistically significant changes between Ts66Yah and Eu mice. When examining age-related alterations, we noted a progressive decline during aging in Ts66Yah mice. Notably, the decline was particularly pronounced from 3 to 6 months and from 3 to 9 months (Figure 6D1). A strong effect of age [F (2, 59) = 9.856, P=0.0002] and genotype [F (1, 59) = 6.872, P=0.0111] was evident. To determine the reduction in autophagy induction in Ts66Yah mice, we analysed the expression of proteins crucially involved in autophagosome nucleation, formation, and maturation. We examined changes in the LC3II/I ratio, a widely used marker of autophagosome formation. The LC3II/I ratio was significantly reduced in 6-month-old Ts66Yah mice compared to Eu mice, although no significant changes were observed at 3 and 9 months in Ts66Yah mice. Significant changes with age were noted for LC3II/I protein within each group [F (2, 60) = 35.52 P<0.0001] (Figure 6G and G1). Further, we examined the expression levels of autophagy-related proteins (ATGs) and, we observed in Ts66Yah mice a significant decrease in ATG7 levels at both 6 and 9 months (Figure 6F). The analysis of ATG5-ATG12 protein levels, did not produced any statistical change (Figure 6E). Finally, we assessed the level of SQSTM1 (p62), a well-established marker of autophagic degradation efficiency. Since SQSTM1 itself is degraded by autolysosomes, its measurement serves as an indirect evaluation of autophagic efficiency, with higher levels indicating reduced autophagic degradation. SQSTM1 protein levels consistently increased at 6 months and 9 months in Ts66Yah mice compared to Eu mice (Figure 6H). The elevation of SQSTM1 protein levels was more pronounced in Ts66Yah mice during aging compared to Eu mice (Figure 6 H1). A significant age and genotype effect was determined (age: [F (2, 57) = 13.34, P<0.0001], genotype: [F (1, 57) = 8.997, P=0.0040]). Building upon our prior findings, which demonstrated early and specific alterations in the UPR and of the PERK branch in DS mice we evaluated the same pathways Ts66Yah hippocampus [16, 17]. Our analysis revealed unchanged levels of BiP/GRP78 protein levels (Figure 7B), while we observed an increase in the active form of PERK (p^T980^/PERK). PERK activation consistently increased at 6 months and 9 months in Ts66Yah mice compared to Eu mice (Figure 7C). The elevation of PERK activation was more pronounced in Ts66Yah mice during aging compared to Eu mice (Figure 7C1). Notably, no significant alterations were detected in PERK protein expression. Genotype significantly influenced alterations in PERK activation [F (1, 59) = 9.312, P=0.0034]. Further, we noted a significant elevation of p^S51^/eIF2α levels only at 6 months of age Ts66Yah respect to the Eu counterparts. According to results obtained with PERK and eIF2α, the downstream target ATF4 demonstrated a substantial increase in 6 and 9 months old Ts66Yah compared to the Eu (Figure 7E and E1). Genotype significantly influenced ATF4 protein expression [F (1, 60) = 19.62, P<0.0001]. In the context of chronic PERK activity, the sustained levels of ATF4 lead to the upregulation of CHOP, which at final stimulates apoptotic responses [28]. Our analysis of nuclear CHOP level showed a substantial increase at 9 months in Ts66Yah mice, while significant variations in CHOP activation within the groups were observed, particularly in Ts66Yah mice, with notable changes occurring from 3 to 6 months and from 6 to 9 months (Figure 7F and F1). A strong influence of age, sex and genotype was detected (age: [F (2, 24) = 5.903, P=0.0082]; sex: [F (1, 24) = 6.514, P=0.0175]; genotype: [F (1, 24) = 4.507, P=0.0443]). Our analysis pointed towards increase autophagy in the Ts66Yah controlled by slightly different mechanism during ageing and suggested a higher level of cellular stress in the brain of DS models.

**Figure 6.**
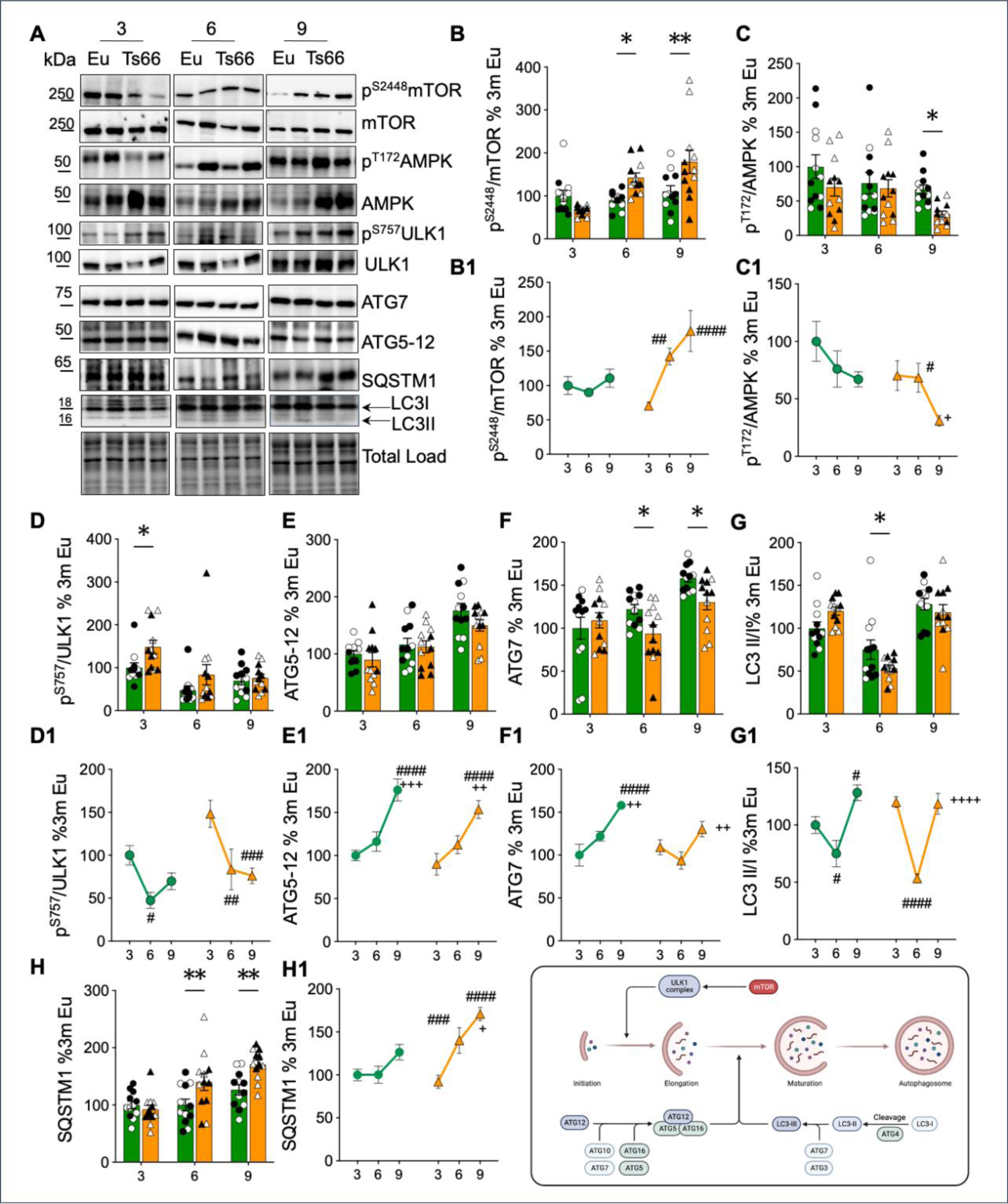
mTOR/AMPK/Autophagy axis evaluation in Ts66Yah hippocampus. Representative western blot images (**A**) and densitometric evaluation of active mTOR (p^S2448^/mTOR) (**B and B1**), AMPK activation (p ^T172^/AMPK) (**C and C1**), ULK1 activation (p ^S757^/ULK1) (**D and D1**), ATG5-12 (**E and E1**), ATG7 (**F and F1**), LC3 II/I (**G and G1**) and SQSTM1 (p62) (**H and H1)** in the hippocampus of Eu and Ts66Yah mice at different ages 3 months (Eu n = 12, Ts66Yah n = 12); 6 months (Eu n = 12, Ts66Yah n = 12); 9 months (Eu n = 12, Ts66Yah n = 12). All densitometric values are given as percentage of Eu at 3 months set as 100%. Data are presented as means ± SEM. Columns were used to show differences among the groups (Eu vs Ts66Yah). Dots were used for Eu and triangle for Ts66Yah. White colour indicates female mice (n =6) and black colour indicate male mice (n =6). Lines were used to show age-associated changes within each group. For columns: *p < 0.05, **p < 0.01, ***p < 0.001 and ****p < 0.0001. For lines: #p < 0.05, ##p < 0.01, ###p < 0.001 and ####p < 0.0001 among 3 months vs 6 and 9 months; +p < 0.05, ++p < 0.01, +++p < 0.001 and ++++p < 0.0001 among 6 months vs 9 months. In (H), scheme relative to the portion of the pathway mentioned in this figure. Created with Biorender.com.

**Figure 7.**
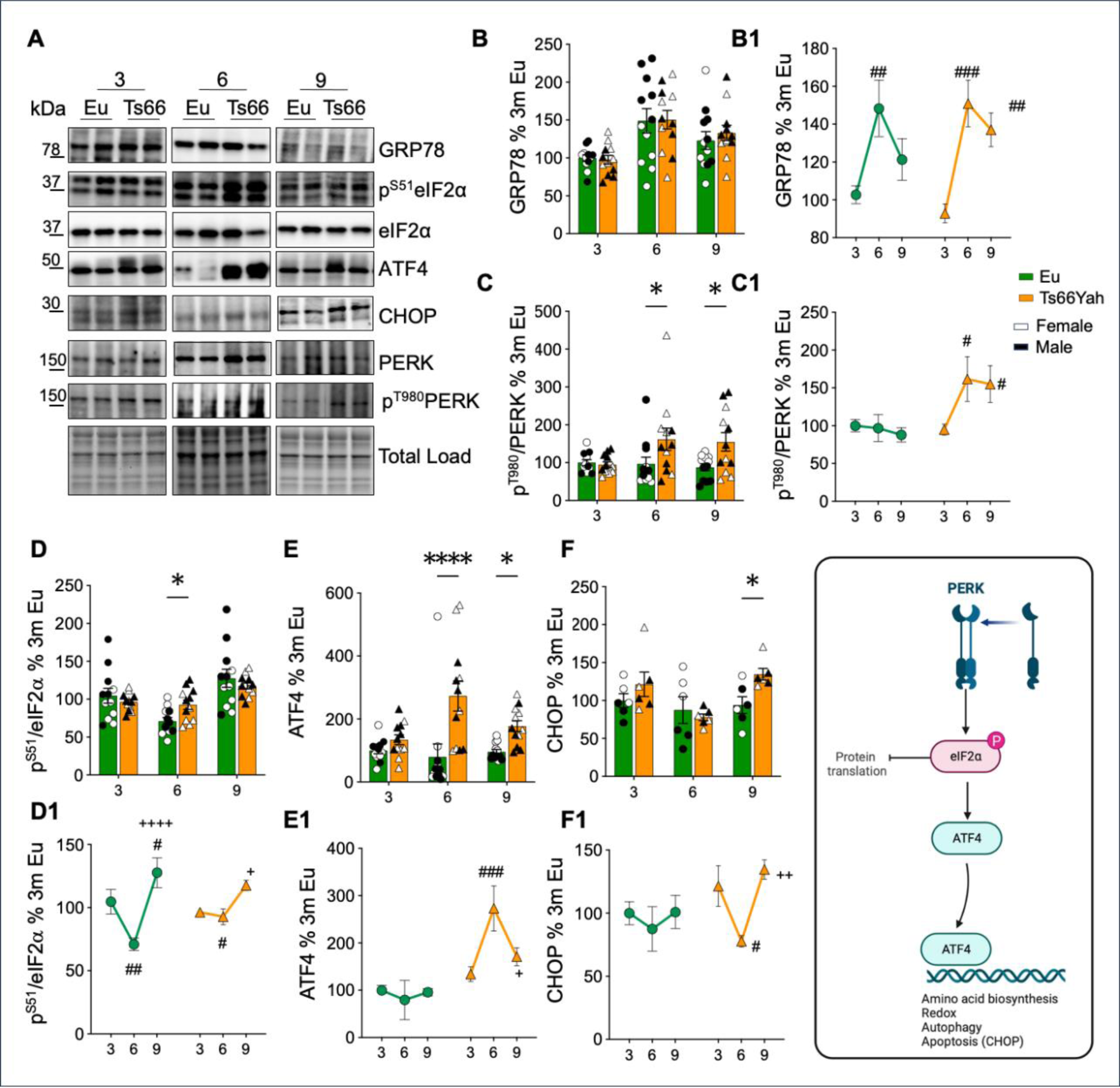
Induction of the PERK pathway in Ts66Yah hippocampus. (**A**) Representative Western blot images and densitometric evaluation of GRP78 (**B and B1**), PERK activation (p^T980^/PERK) (**C and C1**), eIF2α activation (p^S51^/eIF2α) (**D and D1**) and ATF4 (**E and E1**) in the hippocampus of Eu and Ts66Yah mice at different ages 3 months (Eu n = 12, Ts66Yah n = 12); 6 months (Eu n = 12, Ts66Yah n = 12); 9 months (Eu n = 12, Ts66Yah n = 12). Western blot and densitometric analysis of nuclear Chop (**F and F1**) in the above-mentioned age groups: 3 months (Eu n = 6, Ts66Yah n = 6); 6 months (Eu n = 6, Ts66Yah n = 6); 9 months (Eu n = 6, Ts66Yah n = 6). Protein levels were normalized per total protein load. All densitometric values are given as percentage of Eu at 3 months set as 100%. Data are presented as means ± SEM. Columns were used to show differences among the groups (Eu vs Ts66Yah). Dots were used for Eu and triangle for Ts66Yah. White colour indicates female mice and black colour indicate male. Lines were used to show age-associated changes within each group. For columns: *p < 0.05, **p < 0.01, ***p < 0.001 and ****p < 0.0001. For lines: #p < 0.05, ##p < 0.01, ###p < 0.001 and ####p < 0.0001 among 3 months vs 6 and 9 months; +p < 0.05, ++p < 0.01, +++p < 0.001 and ++++p < 0.0001 among 6 months vs 9 months. In (L), scheme relative to the portion of the pathway mentioned in this figure. Created with Biorender.com.

### Increased Apoptosis mechanisms in Ts66Yah mice

TP53/p53 serves as a critical responder to a wide array of cellular stresses capable of causing damage to the cell’s genome. Our findings indicate that p53 expression is altered in Ts66Yah mice. Notably, we observed a significant increase at 3 and 9 months in Ts66Yah mice (Figure 8B and B1), suggesting that trisomy has an impact on p53 expression levels. Furthermore, p53 is the principal transcriptional regulator of *Cdkn1a/p21*, since the gene contains two conserved p53 responsive elements (p53RE) in its promoter. Stress stimuli that elevate p53 activity lead to the increased expression of p21 [30]. Our analysis of p21 protein levels in the nuclear extracts of Ts66Yah mice showed a significant increase at 3, 6 and 9 months (Figure 8C and C1). Genotype significantly influenced p21 activation [genotype: [F (1, 24) = 17.83, P=0.0003]. Furtherly, we analysed BAX, a pro-apoptotic protein that, upon activation, facilitates the release of cytochrome c from mitochondria, ultimately triggering the activation of caspases and the initiation of apoptosis [31]. In our investigation, we noticed elevated BAX protein levels in Ts66Yah compared to Eu mice at 9 months (Figure 8D). In addition is evident for BAX the effect of aging in Ts66Yah mice with a progressive increase from 3 to 9 months (Figure 8 D1). Accordingly, both age [F (2, 60) = 15.95, P<0.0001] and genotype [F (1, 60) = 13.63, P=0.0005] contribute to the variations in BAX levels. The analysis of PARP1 (poly (ADP-ribose) polymerase 1) activation showed a similar with a significant increase in Ts66Yah mice at 9 months old. Caspase 3 and caspase 9 play key roles in the apoptotic cascade, which was reported to be overstimulated in DS phenotype as effect of accelerated aging. As regard, caspase 3 activity and expression, we observed conflicting results among the different age groups of mice. Specifically, the cleaved form of caspase 3, demonstrated in Ts66Yah mice an initial increase at 3 months, followed, however, by a significant decrease at 9 months of age (Figure 8E and E1). Caspase 3 protein expression was instead elevated only at 6 months of age in Ts66Yah mice (Figure 8 F and F1). The alteration of caspase 3 (total and cleaved form) relied mostly on aging and sex effect (aging: [F (2, 60) = 65.33, P<0.0001]; sex [F (2, 60) = 28.17, P<0.0001]). Caspase 9 is demonstrated to be phosphorylated by DYRK1A [32]. Our data show a significant increase in caspase 9 activity, as evidenced by an increase in its cleaved form, in Ts66Yah mice at 3, 6 and 9 months. This increase is likely influenced by genotype [F (1, 60) = 43.31, P<0.0001] (Figure 8H and H1). We also noted a significant increase in caspase 9 protein levels in Ts66Yah mice at 6 and 9 months of age (Figure 8G and G1). These alterations were associated to both genotype [F (1, 60) = 10.88, P=0.0016] and aging [F (2, 60) = 3.462, P=0.0378]. Our results suggest a general induction of the programmed cell death mechanisms as indicated by the increase in proteins linked to the apoptotic machinery and senescence.

**Figure 8.**
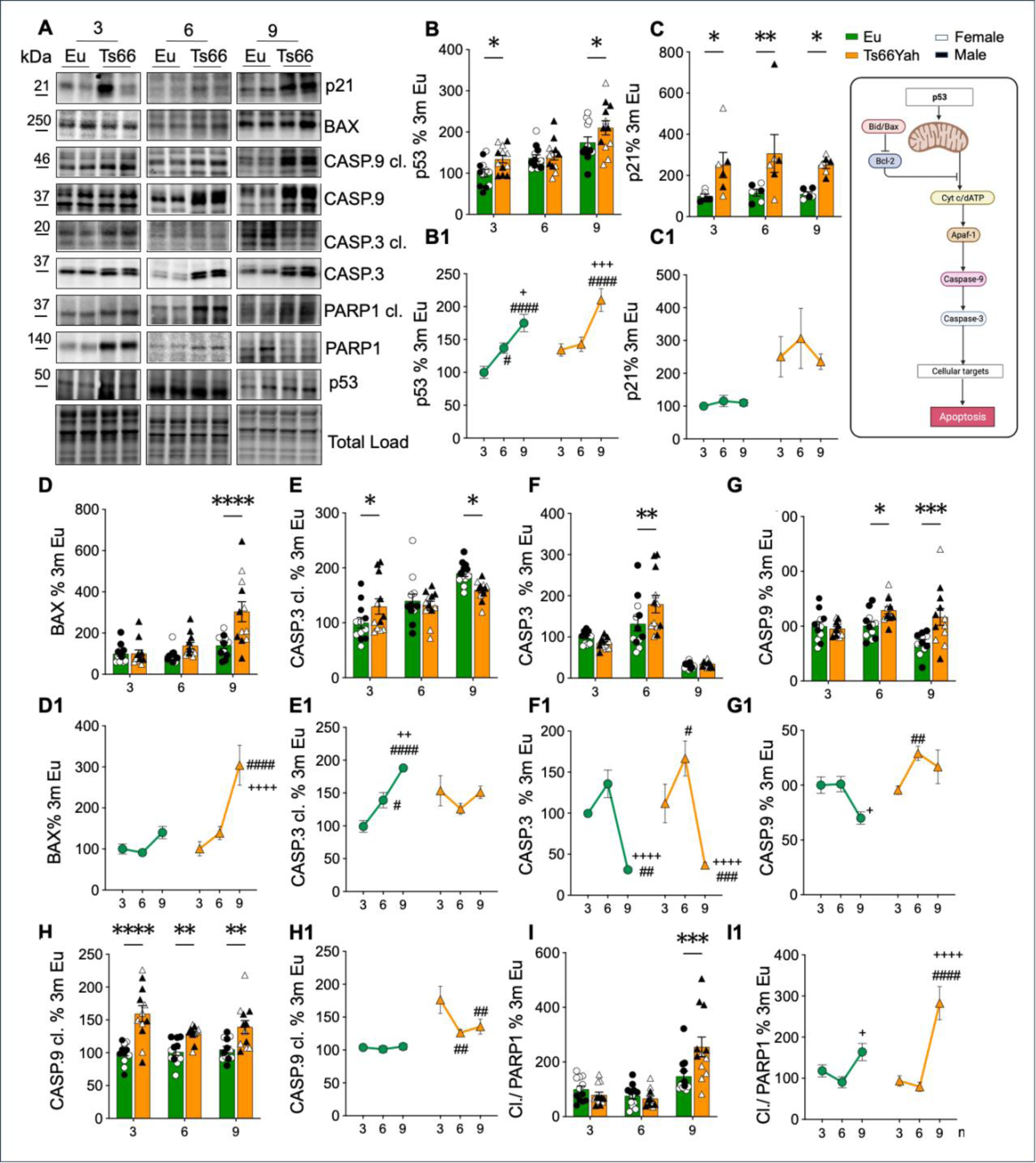
Apoptotic markers induction in Ts66Yah hippocampus. (**A**) Representative Western blot images and densitometric evaluation of p53 (**B and B1**), BAX (**D and D1**), Caspase 3 in its cleaved form and total protein levels (**E, E1**, **F and F1**), Caspase 9 in its cleaved form and total protein levels (**G, G1, H and H1**) and cleaved PARP1/PARP1 ratio (**I and I1**) in the hippocampus of Eu and Ts66Yah mice at different ages [3 months (Eu n = 12, Ts66Yah n = 12); 6 months (Eu n = 12, Ts66Yah n = 12); 9 months (Eu n = 12, Ts66Yah n = 12)]. Western blot images and densitometric evaluation of nuclear p21 (**C and C1**) in the hippocampus of Eu and Ts66Yah mice in the above-mentioned age groups: 3 months (Eu n = 6, Ts66Yah n = 6); 6 months (Eu n = 6, Ts66Yah n = 6); 9 months (Eu n = 6, Ts66Yah n = 6). Protein levels were normalized per total protein load. All densitometric values are given as percentage of Eu at 3 months set as 100%. Data are presented as means ± SEM. Columns were used to show differences among the groups (Eu vs Ts66Yah). Dots were used for Eu and triangle for Ts66Yah. White colour indicates female mice and black colour indicate male mice. Lines were used to show age-associated changes within each group. For columns: *p < 0.05, **p < 0.01, ***p < 0.001 and ****p < 0.0001. For lines: #p < 0.05, ##p < 0.01, ###p < 0.001 and ####p < 0.0001 among 3 months vs 6 and 9 months; +p < 0.05, ++p < 0.01, +++p < 0.001 and ++++p < 0.0001 among 6 months vs 9 months. In (C), scheme relative to the portion of the pathway mentioned in this figure. Created with Biorender.com

### Alteration of synaptic plasticity markers in Ts66Yah mice

To elucidate the cognitive and learning dysfunctions highlighted in the behavioural tests, we delved into the changes in two crucial synaptic proteins: syntaxin-1 (pre-synaptic) and postsynaptic density protein 95 (PSD95). We also wondered if aging could have more influence on some synaptic markers in the DS model. Syntaxin-1 is a neuronal plasma membrane protein involved in vesicle trafficking, docking, and fusion, playing a pivotal role in neurotransmitter release. On the other hand, PSD95, a membrane-associated guanylate kinase, serves as the major scaffolding protein in the excitatory postsynaptic density (PSD) and plays a significant role in regulating synaptic strength [24]. In Ts66Yah mice, we observed an unexpected and significant increase in Syntaxin-1 levels at 3 months of age compared to Eu mice. However, as mice aged, protein levels exhibited a significant reduction at 6 months and although not statistically significant, there was a trend towards further reduction in 9-month-old Ts66Yah mice (Figure 9B). Additionally, age-related changes in Syntaxin-1 within each group highlighted a decrease in Ts66Yah in the transition from young to adult mice (aging:[F (2, 60) = 5.071, P=0.0092]; Figure 9B1). As observed for Syntaxin-1, PSD95 protein levels exhibited a significant increase at 3 months in Ts66Yah mice compared to the Eu group. However, in mature adult mice (9 months) PSD95 levels decreased significantly (Figure 9C). The evaluation of age-related changes within each group revealed a strikingly different trajectory between Eu and Ts66Yah mice, underscoring an aberrant expression of PSD95 protein in the early stages of Ts66 mice, which dramatically declined in later phases (Figure 9C1). A strong influence of age [F (1, 60) = 34.49, P<0.0001] was detected. Moreover, we evaluated the levels of brain-derived neurotrophic factor (BDNF), a neurotrophin implicated in learning and memory functions. BDNF protein levels were significantly increased in Ts66Yah mice at 6 months, but then declined at 9 months suggesting that inadequate neurotrophic support from BDNF may contribute to neurodegeneration and cognitive decay (Figure 9D and D1).

**Figure 9.**
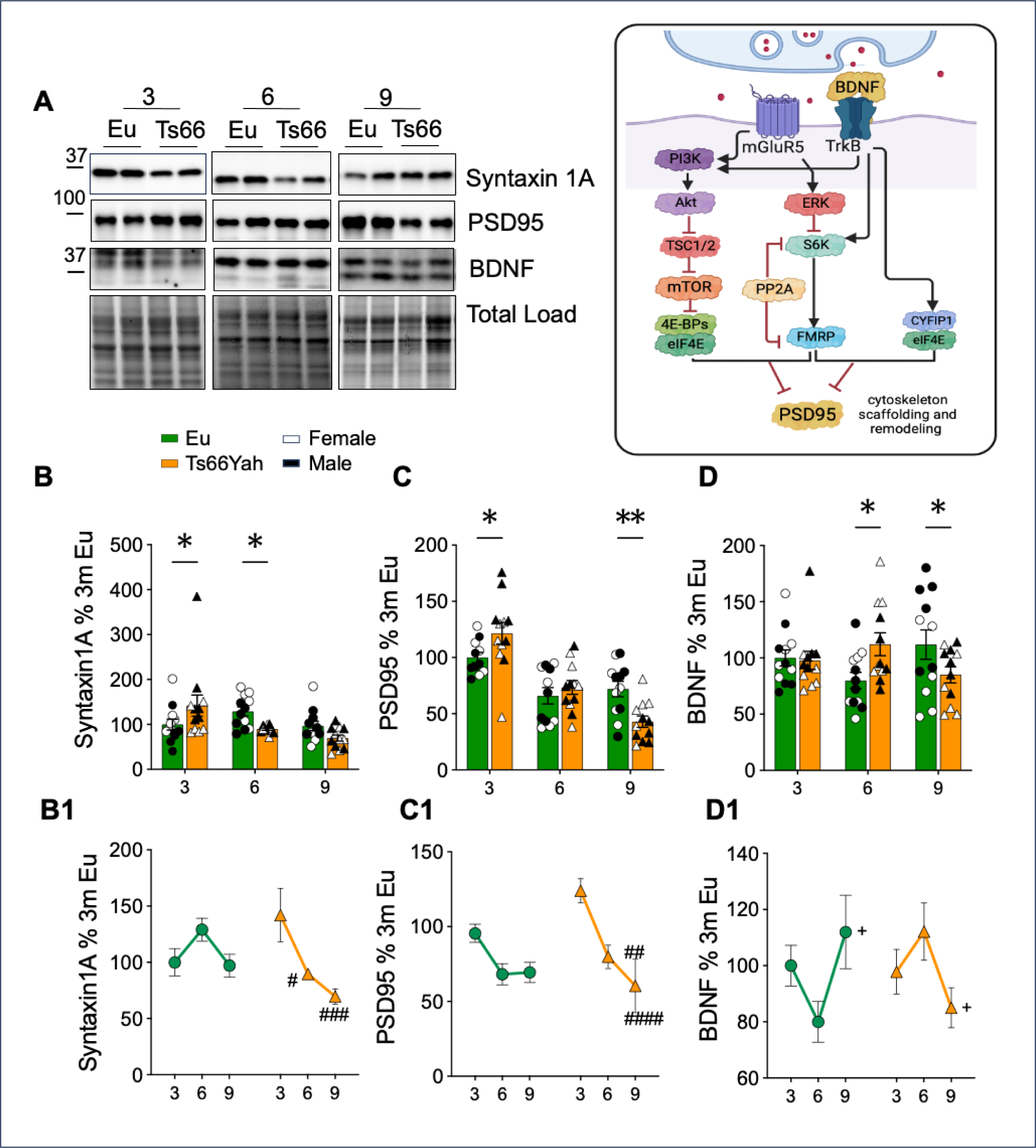
Markers of synaptic plasticity in Ts66Yah hippocampus. Syntaxin1, PSD95 and BDNF protein levels were evaluated in the hippocampus of Eu and Ts66Yah mice at different ages 3 months (Eu n = 12, Ts66Yah n = 12); 6 months (Eu n = 12, Ts66Yah n = 12); 9 months (Eu n = 12, Ts66Yah n = 12). (**A**) Representative Western blot images and densitometric evaluation of (**B and B1**) syntaxin-1, (**C and C1**) PSD95 and (**D and D1**) BDNF. All densitometric values are given as percentage of Eu at 3 months set as 100%. Data are presented as means ± SEM. Columns were used to show differences among the groups (Eu vs Ts66Yah). Dots were used for Eu and triangle for Ts66Yah. White colour indicates female mice (n =6) and black colour indicate male mice (n =6). Lines were used to show age-associated changes within each group. For columns: *p < 0.05, **p < 0.01, ***p < 0.001 and ****p < 0.0001. For lines: #p < 0.05, ##p < 0.01, ###p < 0.001 and ####p < 0.0001 among 3 months vs 6 and 9 months; +p < 0.05, ++p < 0.01, +++p < 0.001 and ++++p < 0.0001 among 6 months vs 9 months. In (A), scheme relative to the portion of the pathway mentioned in this figure, with highlighted in yellow proteins evaluated by Western blot analysis. In (A), scheme relative to the portion of the pathway mentioned in this figure. Created with Biorender.com.

## DISCUSSION AND CONCLUSIONS

The genetic validity of available murine models for the study of DS phenotype has been a topic of strong debate in the last decade [3, 33–36]. To improve the representation of DS human genotype and phenotype in mice, Herault and colleagues developed new line named Ts66Yah, derived from the Ts65Dn lineage but no longer carrying the duplicated centromeric part of Mmu17, [20]. Recent studies compared the phenotype of the Ts66Yah and Ts65Dn mouse models of DS in terms of learning and behaviour, craniofacial and brain morphologies, and gene expression, confirming the interference of *Scaf8-Pde10a* region trisomy in Ts65Dn. Overall, Ts66Yah showed fewer dysregulated pathways in the EC and similar learning and memory behavioural deficits [20]. Behavioural analysis performed in this longitudinal study on 3 and 9 months old Ts66Yah mice support previous findings along ageing and extend the relevance of challenging more the mice to detect a phenotype in spatial memory. For the spatial memory, decreasing the learning trials induced memory deficits in young and older individuals, compared to what has been previously described in the young [20]. Similar changes in the training could influence also other behavioural outcomes in DS models. Having add one additional training sessions for the mutant mice to investigate the familiar object make them performed better in [21] than in the original publication where it was tested in two independent labs [20]. This phenomenon is probably reminiscent in human when specific education program is available, people with DS performed better. More interestingly this report highlighted different ageing effect in the Ts66Yah model mainly affecting the spatial memory while the ageing effect on the two other behaviour domains investigated for object memory or anxiety-related behaviour, followed the same trajectories in Ts66Yah and Eu control genotypes.

In order to decipher the age-related proteome changes in the hippocampus from Ts66Yah compared to euploid mice, we investigated how the overexpression of protein products from trisomic genes may influence specific neuronal pathways whose dysfunction is associated with the neurodegenerative process. Indeed, our analysis foresees to evaluate the proteostasis network and related stress responses, metabolic processes, cell death mechanisms and synaptic transmission, whose defects in induction, regulation or function represents well-established fingerprints of DS brain phenotype [17, 24, 37, 38]. As expected, the analysis of protein products from genes mapping in triplicated regions demonstrated increased expression levels in Ts66Yah mice (Figure 10). Particularly, the investigation of APP, DYRK1A, SOD1 and BACH1 support the early genotype-related development of hippocampal degenerative signatures, as observed in Ts65Dn mice and in other DS models. The overexpression of APP was demonstrated to be sufficient to induce the increase in the production of Aβ peptides, which represent crucial triggers of neurodegeneration despite the lack of senile plaques formation in mice [39, 40]. As well, the overexpression and hyperphosphorylation of TAU occurs early in Ts66Yah hippocampus and it was demonstrated to promote microtubular dysfunction and neuronal degeneration associated with Alzheimer-like pathology [26, 41]. The noxious relationship between APP and TAU is also supported by a network analysis using the Ingenuity Pathway Analysis software (Fig. 10 C). Most likely trisomy-related APP overexpression may indirectly dysregulate and trigger the early increase of tau levels. Besides, the overexpression of DYRK1A, which was demonstrated to target and phosphorylate TAU also on Ser202, is in line with the observed increase of p-tau levels in Ts66Yah and confirm the predisposition of this DS model in developing AD-signatures early in life [42–44].

**Figure 10.**
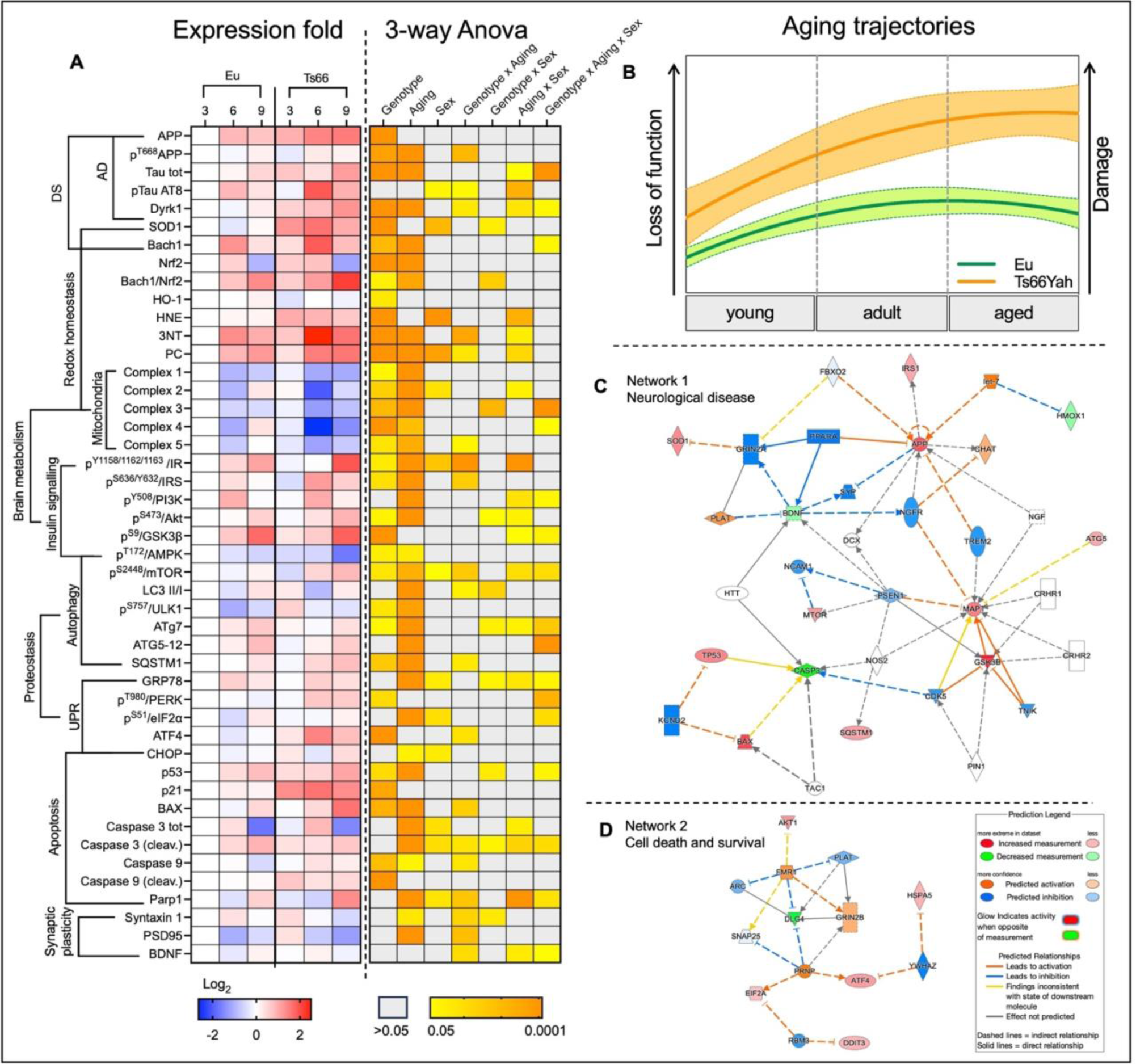
Overview of data expression differences and molecular networking. **(A)** The Heat-map on the left shows changes observed for the proteins analysed in the study. The heatmap shows the relative abundance (Log_2_ of expression ratio) for all the proteins analysed grouped according to the increase of aging in the different genotype groups. The proteins are clustered according to their signalling pathway. The white square indicates Euploids mice at 3 months of age set as 0 and used as normalization group. The heat map on the right indicates the influence of genotype, aging and their interaction for all the molecular markers analysed **(B).** Aging trajectory in Ts66Yah (orange) and Eu (green) animals built from data collected in the present study. **(C)** Network analysis concerning the protein differentially expressed/activated between Ts66Yah and Eu mice. Network were built using Ingenuity Pathway Analysis (Qiagen). **(C)** Network 1 ”Neurological Disease” 35 proteins components **(D)** Network 2 ”Cell Death and Survival” 14 proteins components.

**Figure 11.**
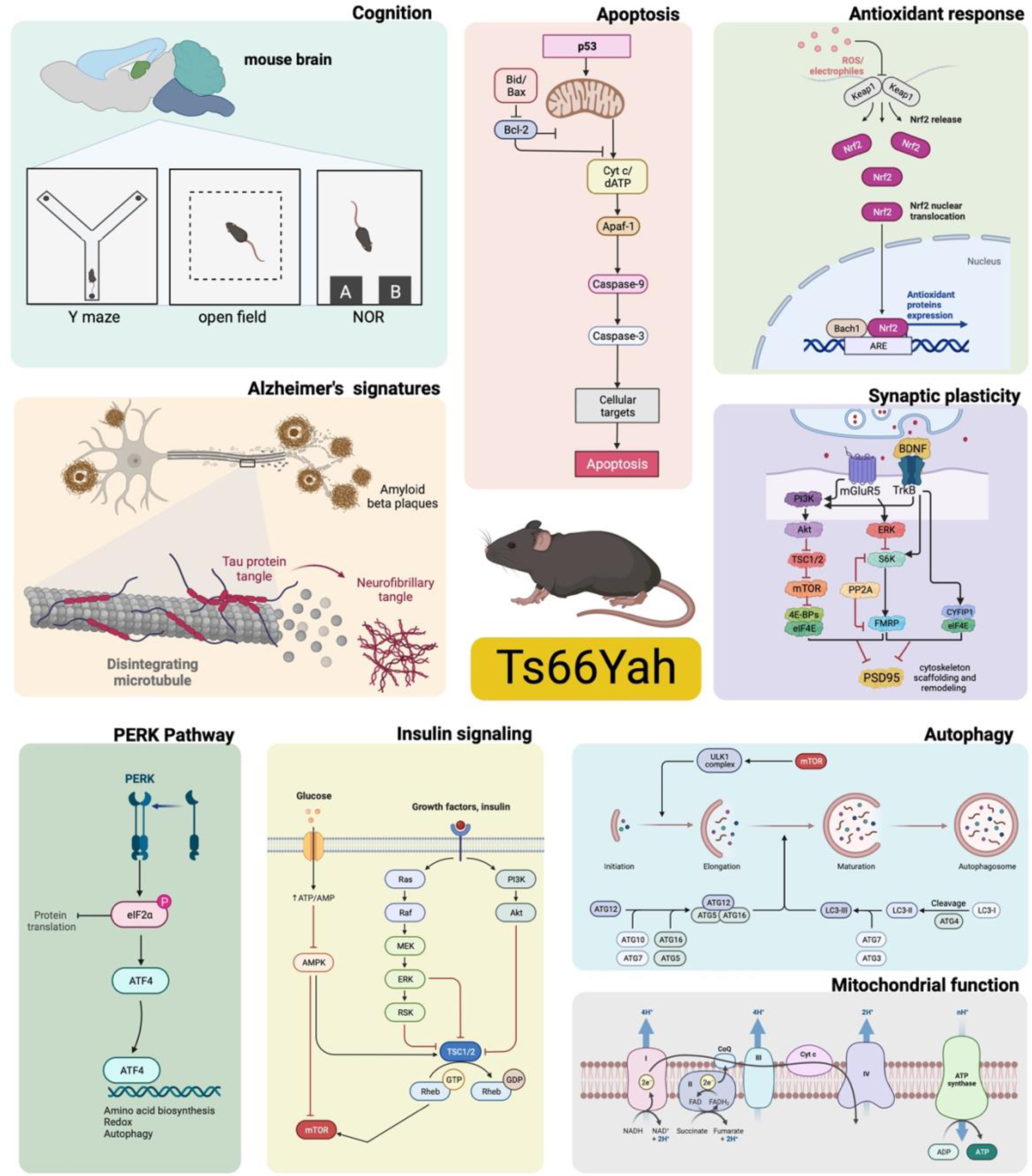
Schematic overview of molecular mechanisms and signalling pathways investigated in Ts66Yah mouse model. Created with Biorender.com.

The premature alteration of redox homeostasis, demonstrated in Ts66Yah mice by increased oxidative damage to proteins, is a pathological feature of the DS brain actively contributing to the degenerative process [28, 37, 45]. The triplication of *SOD1* and *BACH1* genes has been associated in DS with the reduction of brain antioxidant capacity impinging in the overproduction of H_2_O_2_ and in the depletion of NRF2 antioxidant response. Intriguingly, while SOD1 overexpression shows a robust increase since early age in Ts66Yah mice, the alteration of BACH1/NRF2 axis primes in young animals but rises and consolidates in older mice as effect of both genotype and aging. These data confirm the potential toxicity of the unbalanced levels of endogenous antioxidant enzymes, as postulated in previous studies [27, 28]. Furthermore, the depletion of NRF2-rmediated antioxidant response, associated with the overexpression of BACH1, appears to be a common mechanism of neurodegeneration also observed in AD neuropathology [17]. The loss of redox homeostasis is often tied with the alteration of the mitochondrial OXPHOS machinery. Indeed, the proper functionality of mitochondrial ETC is fundamental for energy production and regulates brain development and homeostasis, since many processes, including stress response mechanisms, relies on ATP consumption. On the contrary, defective mitochondria become the first source of ROS that spread to other cell compartments and damage biological molecules particularly in the presence of reduced antioxidant capacity, as observed in Ts66Yah mice. Mitochondrial impairment is one of the earliest events observed in DS neurodegeneration [46, 47] and mitochondrial dysfunction has been extensively reported in Ts65Dn mice [24, 48]. Our data on the Ts66Yah model supports previous findings, by showing the dysfunction of complex I expression in 3 months-old animals, which suggest an early compromission of mitochondrial functionality, as a direct effect of genotype. However, a significant reduction of complexes I, II, IV and V is observed in old mice under the additional influence of aging.

Besides mitochondrial defects, the alteration of energy metabolism reported in DS is further linked with the development of brain insulin resistance (BIR) [24, 49, 50]. The failure of insulin to stimulate its signalling cascade may, on one side alter glucose catabolism, thus exacerbating the energetic decay reported in the brain, and on the other side disturb brain development [51]. Indeed, insulin serves also as a trophic factor, which regulates neural stem cells proliferation, differentiation and survival. Disturbances of the insulin signalling are indexed by the uncoupling between IR and IRS1 activation in Ts66Yah animals at 6 and 9 months of age. Such uncoupling is confirmed by the lack of PI3K induction, while AKT independently is increasingly active only at 9 months as also previously seen in DS humans and mice [24, 50]. Intriguingly, no differences were observed for GSK3β activation between Ts66Yah and Eu mice at different ages, while an age-dependent increase of GSK3β inhibitory state is observed in both Ts66Yah and Eu mice aged from 3 to 9 months. Downstream of AKT the activation of mTORC1 represents a key mechanism fine tuning proteostasis during brain development and aging [52]. A dysregulation of mTOR activity occurs during pathological condition associated with metabolic stressors, such as BIR, and with the accumulation of toxic aggregates (e.g., Aβ). Aberrant mTORC1 induction was observed in DS humans prior and after AD development and in the brain of different mouse models of the disease [15, 38, 53]. Data collected on Ts66Yah are in line with previous findings reporting increased mTORC1 activation in 6- and 9-months old transgenic animals as effect of both genotype and aging. Active mTORC1 exerts an inhibitory effect on autophagy induction by reducing the initial phases such as autophagosome formation and maturation. However, mTORC1 influence on autophagy is antagonized by AMPK, a major regulator of the autophagic machinery, which is activated by high AMP/ATP ratio [51]. Phosphorylated AMPK induces autophagy by ULK1 activation and by mTORC1 inhibition via phosphorylation of Raptor. Both AMPK and mTOR also control cell growth and metabolism, coupling autophagy to these processes. According to mTORC1 data, AMPK demonstrates a trend of decreased phosphorylation in Ts66Yah mice that becomes significant in the 9 months old group. The opposite behaviour among mTORC1 and AMPK promote the reduced induction of autophagy demonstrated by decreased LC3 II/I ratio, reduced levels of ATG7 and increased levels of SQSTM1 (p62) in Ts66Yah mice at 6 and 9 months of age. Reduced autophagy contributes to the failure of protein degradation mechanisms with consequent accumulation of protein damage and toxic aggregates, as observed in DS [54–56]. The importance of mTORC1/autophagy axis in the maintenance of brain homeostasis and intellect is furtherly strengthened by data on rapamycin treatment collected in Ts65Dn, where the amelioration of hippocampal neuropathology, improved cognitive performances and reduced proteotoxicity [38, 57]. In addition, results collected on Ts66Yah mice agree with studies showing the occurrence of endosomal pathology in DS human and mice as results of gene triplication [58, 59]. Compelling studies in DS demonstrated that, beyond the failure of protein degradation pathways, proteostasis is perturbed by the aberrant induction of protein re-folding mechanisms, which under chronic stress conditions (e.g., ER stress) becomes detrimental for brain cells [28, 60]. An overactivation of PERK- and PKR-related responses was observed in DS mice leading to a general repression of protein translation as effect of eIF2α induction [16, 17, 61]. Data collected in Ts66Yah, confirm previous findings and support the overactivation of the PERK pathway in 6- and 9-months old animals and its uncoupling with the induction of NRF2 response [17]. The induction of the PERK branch of the UPR lead to eIF2α phosphorylation and ATF4 increased expression at 6 and 9 months of age. Based on the extent of its activation, ATF4 plays a critical role in cell adaptation or in the activation of apoptosis. The sustained ATF4 levels reported in Ts66Yah lead to the upregulation of nuclear CHOP in 9 months old animals, which in turn promotes apoptosis by enhancing expression of DR5 and tribbles-related protein 3. However, the induction of the apoptotic machinery through CHOP is paralleled, in DS mice, by the activation of p53 signal. p53 play a crucial role in regulating the transcription of a wide set of genes involved in cell cycle arrest, senescence, antioxidant system or apoptosis in response to various stress signals. Findings in DS reported the uncontrolled increase of p53 induction that may have deleterious effects leading to an “peculiar” pro-apoptotic phenotype [31]. Data on Ts66Yah support the early genotype-dependent induction of p53 signal, as demonstrated by the increased expression of p53, p21 (downstream signal), caspase 3 (cleaved form) and caspase 9 at 3 months of age. Such activation of apoptosis machinery is maintained and furtherly exacerbated in old animals, as demonstrated by BAX and PARP1 expression, also supporting the influence of aging in the altered induction of cell-death responses. Finally, to tie molecular alterations and cognitive decline observed in Ts66Yah mice we analysed the expression of the pre-synaptic protein syntaxin 1, post-synaptic protein PSD-95 and of the neurotrophin BDNF [24]. We observed the early increased expression for syntaxin 1 and PSD-95, which suggest the activation of mechanisms to cope brain alterations associated with genotype and maintain synaptic plasticity. However, such effect is lost in aged mice where decreased levels of PSD-95 and BDNF parallel the reduced cognition described above.

This research presents a set of data exploring both mice cognition and various molecular pathways associated with changes in the DS phenotype, likely playing a role in the progression of neurodegeneration. On the hole, results collected in Ts66Yah mice report an early and consistent effect of the genotype, which intensifies under the influence of aging. Besides, sex differences did not exhibit pronounced effects in this DS murine model. Distinctly, cognitive testing underscored different aging effects in the Ts66Yah model, primarily impacting spatial memory, while object memory and anxiety-related behavior followed similar aging trajectories in both Ts66Yah and Eu genotypes. Alongside, the analysis of molecular markers revealed altered expression/activation that was already significant in young Ts66Yah mice. This evidence indicates that genotype-related changes manifest early, affecting cellular mechanisms at various levels, thereby influencing brain development, and fostering a state of premature aging. Remarkably, our biochemical observations reveal that gene triplication leads to cognitive decline and brain pathology, impacting both metabolic pathways and stress-related responses. Metabolic alterations, on one side, can contribute to energy failure and the production of ROS, disrupting brain homeostasis. Conversely, the diminished activation of stress responses hinders the detoxification of brain cells from proteotoxic insults, leading to the build-up of protein aggregates and exacerbating brain damage. Hence, the interplay between these detrimental events may establish a vicious cycle, impairing brain maturation initially, and promoting neurodegeneration later in life. In line with this, Ts66Yah and Eu mice exhibit a similar aging trajectory for the studied molecular markers, yet they diverge at the outset due to the prime impact of genotype (Figure 10 B). As a result, the data from Ts66Yah mice indicate that molecular changes in the brain initiate due to genotype differences and subsequently progress with aging, highlighting a synergistic effect between these two factors in shaping the DS phenotype. Moreover, we propose that Ts66Yah mice recapitulate the majority of cognitive and molecular alterations associated with genotype and aging observed in previous partial DS mouse models. Importantly, these findings offer novel and consolidated insights, devoid from genomic bias, into trisomy-driven processes that may contribute to defects in brain maturity and subsequently progress to neurodegeneration. Additionally, the results from Ts66Yah mice, by identifying both data commonalities and differences, contribute to bridging the knowledge gap in understanding the complexity of DS phenotypes. This may, in turn, refine the scientific significance of studies conducted on Ts65Dn and on related DS models.

## Acknowledgments

N/A

## Author contribution

Conceptualization: FDD, MP, YH

Data Curation: MRB, CL, VN, FDD, YH, FP, EB

Formal Analysis: MRB, FL, FP, CS

Funding Acquisition: FDD, MP, YH

Investigation: MRB, CL, VN, ML, FDD, YH, AT, FP, CS

Methodology: MRB, CL, FDD, VN, AT, FP, CS, AD

Project Administration: FDD, MP, YH

Supervision: FDD, MP, YH

Validation: MRB, CL, FDD, MP, YH

Visualization: MRB, CL, FDD, VN, YH

Writing – Original Draft Preparation: CL, FDD, MP, YH

Writing – Review & Editing All

