## Supplementary material for "Cognitive and molecular characterization of the Ts66Yah murine model of Down syndrome: deepening on hippocampal changes associated with genotype and aging": Supplemetary table 1

**Supplementary table 1:** List of antibodies used in this study

| **Antibodies** | **Brand** | **Diluition** | **Catologue** |
| --- | --- | --- | --- |
| IRβ | Cell signaling tecnology | 1:1000 | #3030 |
| Phospho (Tyr1158/1162/1163)-Irβ | Genetex | 1:1000 | GTX25681 |
| anti-IRS1 | Cell signaling tecnology | 1:1000 | #3407 |
| Phospho (Ser636)-IRS1 | Santa Cruz Biotechnology | 1:500 | sc-33957 |
| Phospho (Tyr632)-IRS1 | Santa Cruz Biotechnology | 1:500 | sc-17196 |
| 3NT | Sigma Aldrich | 1:1000 | N5538 |
| HNE | Novus Biologicals | 1:2000 | Nb-10063093 |
| anti-DNP | Merck Millipore | 1:5000 | #90451 |
| Tau total | Byorbit | 1:500 | orb-46243 |
| phosphoTau (Ser202, Thr205) AT8 | Invitrogen | 1:1000 | MN1020 |
| Atg7 | Cell signaling Technology | 1:1000 | D12B11 |
| Atg5-Atg12 | Invitrogen | 1:1000 | 702433 |
| p-AKT (ser 473) | Cell signaling Technology | 1:1000 | #93H12 |
| AKT | Biorad Laboratories | 1:1000 | vma00253K |
| GSK3b (C-terminal) | Abcam | 1:1000 | ab93926 |
| Phospho (Ser9)-GSK-3β | Cell signaling tecnology | 1:1000 | #5558 |
| total OXPHOS | Abcam | 1:5000 | ab110411 |
| Complex 1 (NDUFB8) | Novus Biological | 1:1000 | NBP2-7558 |
| ATF4 | Santa Cruz Biotechnology | 1:500 | sc-390063 |
| APP | Sigma Aldrich | 1:10000 | A8717 |
| phospho-APP | Invitrogen | 1:1000 | PA5-97328 |
| CHOP | Invitrogen | 1:1000 | MA1-250 |
| SOD1 | Santa Cruz Biotechnology | 1:500 | sc-271014 |
| BACH-1 | Byorbit | 1:500 | orb4401 |
| Nrf2 | Gene tex | 1:1000 | GTX 103322 |
| Syntaxin-1 | Gene tex | 1:1000 | GTX113559 |
| PSD95 | Abcam | 1:1000 | ab18258 |
| BDNF | Santa Cruz Biotechnology | 1:1000 | sc-546 |
| phospho-mTOR (Ser2448) | Cell signaling Technology | 1:1000 | 5536S |
| mTOR | Cell signaling Technology | 1:1000 | #4517 |
| DYRK1A D30C10 | Cell signaling Technology | 1:1000 | #8765 |
| p21 | Sigma Aldrich | 1:1000 | 05-655 |
| Pol II (F-12) | Santa Cruz Biotechnology | 1:1000 | sc-55492 |
| SQSTM1/p62 | Gene tex | 1:1000 | GTX100685 |
| ULK 1 | Cell signaling Technology | 1:1000 | 8054S |
| phospho-ULK1 (ser757) | Invitrogen | 1:1000 | PA5-105130 |
| PI3-kinase p85α (N-18) | Santa Cruz Biotechnology | 1:500 | sc-31969 |
| [p-PI 3-kinase p85α (Tyr 508)](https://datasheets.scbt.com/sc-12929.pdf) | Santa Cruz Biotechnology | 1:500 | sc-12929 |
| p53 | Santa Cruz Biotechnology | 1:1000 | SC-393031 |
| Cleaved Caspase-3 | Cell signaling Technology | 1:1000 | #9915 |
| Caspase-3 |  | 1:1000 | #9915 |
| Cleaved Caspase-9 |  | 1:1000 | #9915 |
| Caspase-9 |  | 1:1000 | #9915 |
| Cleaved PARP |  | 1:1000 | #9915 |
| PARP |  | 1:1000 | #9915 |
| Bax | Santa Cruz Biotechnology | 1:1000 | sc-23959 |
| LC3B | Novus Biologicals | 1:1000 | NB-1002220 |
| ampk | Santa Cruz Biotechnology | 1:500 | sc-74461 |
| [Phospho-AMPKα (Thr172) (40H9)](https://www.cellsignal.com/products/primary-antibodies/phospho-ampka-thr172-40h9-rabbit-mab/2535) | Cell signaling Technology | 1:1000 | 2535S |
| perk | Santa Cruz Biotechnology | 1:1000 | sc-377400 |
| [Phospho-PERK (Thr980)](https://www.thermofisher.com/antibody/product/Phospho-PERK-Thr980-Antibody-Polyclonal/BS-3330R) | Bioss | 1:1000 | bs-3330R |
| eif2a | Cell signaling Technology | 1:1000 | 5324S |
| [Phospho-eIF2α (Ser51)](https://www.cellsignal.com/products/primary-antibodies/phospho-eif2a-ser51-d9g8-xp-rabbit-mab/3398) | Cell signaling Technology | 1:1000 | 3398S |
| HO-1 | Enzo Life sciences | 1:1000 | ADISPA 896F |
| Grp78 | Santa Cruz Biotechnology | 1:1000 | sc-376768 |
