## Supplementary figures 1-5 for "Cognitive and molecular characterization of the Ts66Yah murine model of Down syndrome: deepening on hippocampal changes associated with genotype and aging"

**Supplementary Fugures**

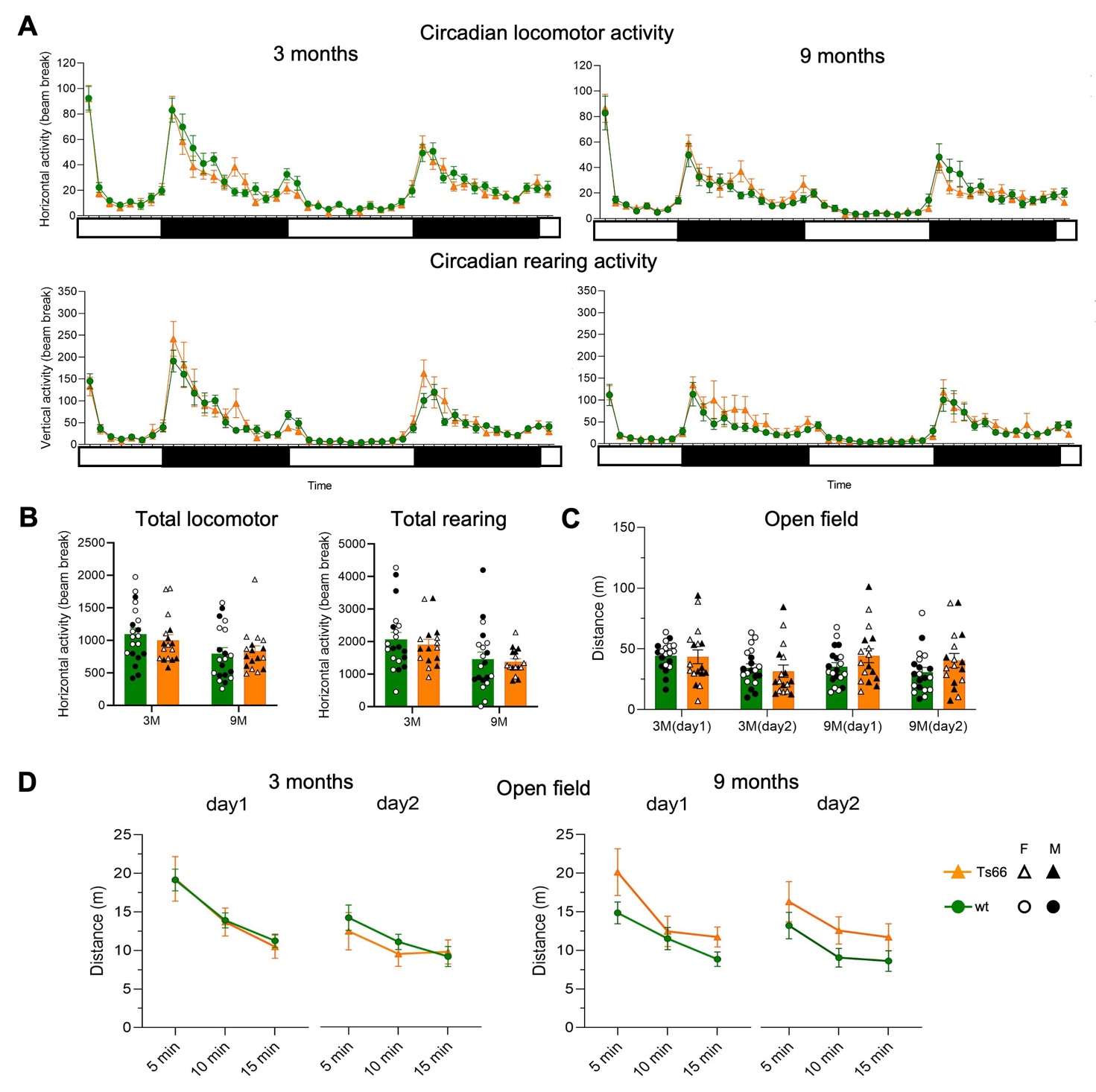

**Supplementary Figure 1. Motor activity is not affected in Ts66Yah mice.** No difference in horizontal (A) and vertical (B) activity was detected in Ts66mice comparing to control littermates during circadian activity (CA) at 3 and 9 months of age (3M and 9M). There was no significant difference between Ts66Yah mice and control littermates in distance travelled or in time spent in center of the arena in the open field (OF) test at 3 and 9 months (C). Distance travelled during the open field session at 3 and 9 months showing similar reduced exploration along time with 10min intervals (D).

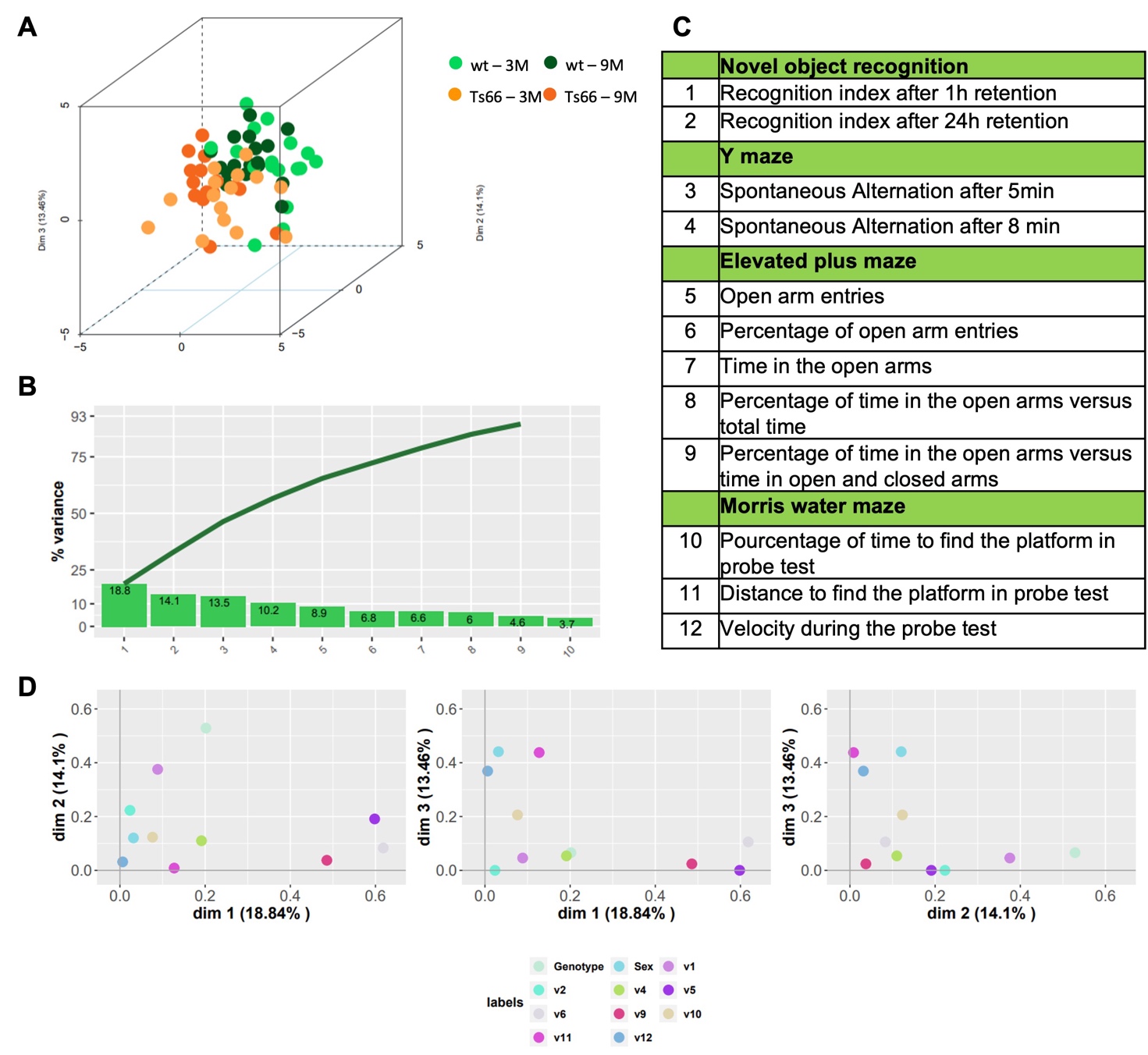

**Supplementary Figure 2.** Multifactorial analysis of the Ts66Yah mouse compared to control littermates (Eu) with 11 non-correlated out of 12 behavioural variables assessed at 3 and 9 months. (A) 3D representation showing the relative position of each individual used in the analysis. (B) Contribution of each dimension to the global percentage of variance. (C) List of 12 variables used in the analysis, (D) Contribution of each behavioural variable to the first three dimensions.

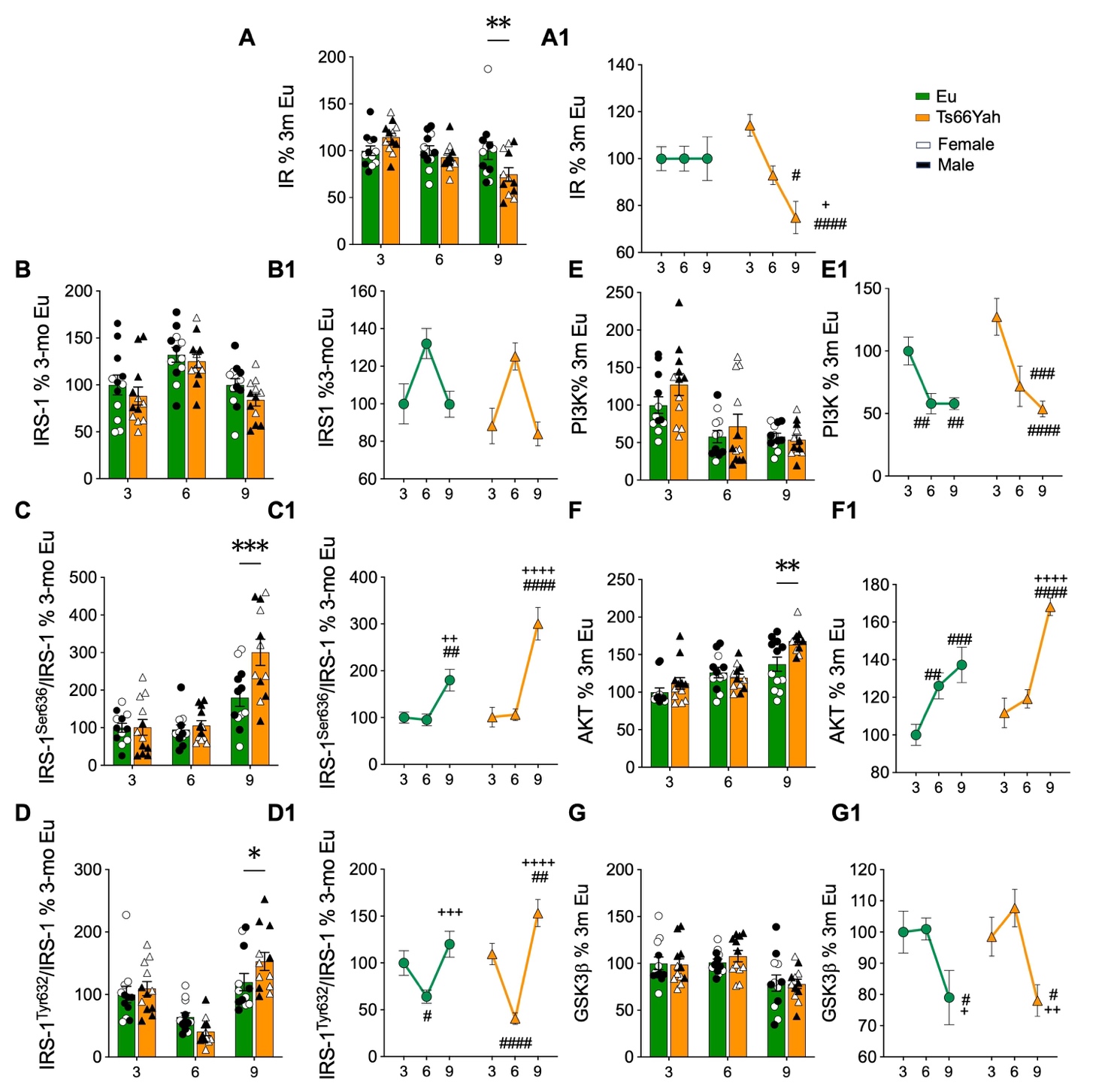

**Supplementary Figure 3.** Densitometric evaluation of IR (**A and A1**), IRS-1(**B and B1**), IRS-1^Ser636^/IRS1 (**C and C1**), IRS-1^Tyr632^/IRS1 (**D and D1**), PI3K (**E and E1**), AKT **(F and F1)** and GSK3B **(G and G1)** in the hippocampus of Eu and Ts66Yah mice at different ages 3 months (Eu n = 12, Ts66Yah n = 12); 6 months (Eu n = 12, Ts66Yah n = 12); 9 months (Eu n = 12, Ts66Yah n = 12). Protein levels were normalized per total protein load. All densitometric values are given as percentage of Eu at 3 months set as 100%. Data are presented as means ± SEM. Columns were used to show differences among the groups (Eu vs Ts66Yah). Lines were used to show age-associated changes within each group. For columns: *p < 0.05, **p < 0.01, ***p < 0.001 and ****p < 0.0001. For lines: #p < 0.05, ##p < 0.01, ###p < 0.001 and ####p < 0.0001 among 3 months vs 6 and 9 months; +p < 0.05, ++p < 0.01, +++p < 0.001 and ++++p < 0.0001 among 6 months vs 9 months.

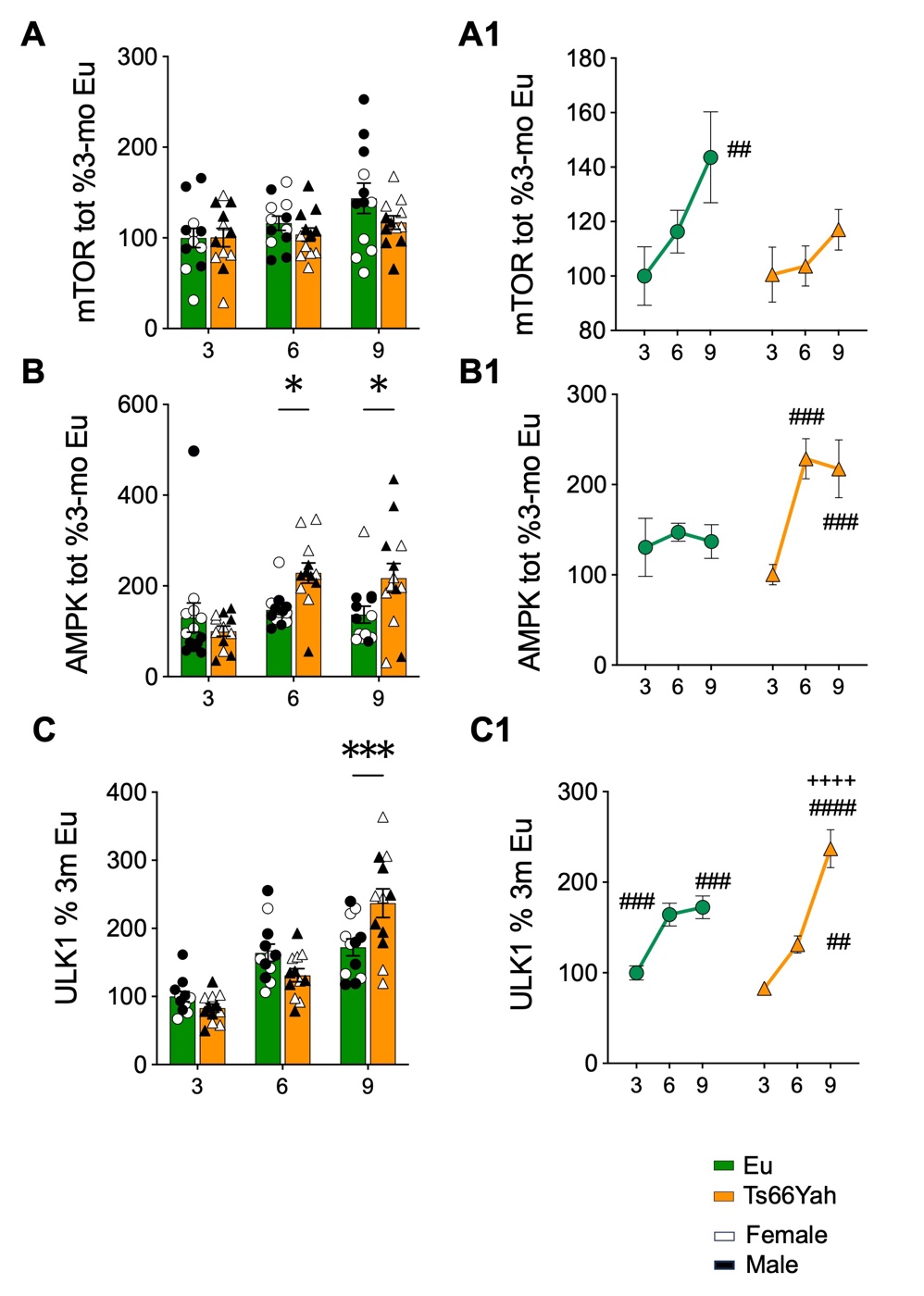

**Supplementary figure 4.** Densitometric evaluation of mTOR (**A and A1**), AMPK (**B and B1**) and ULK1 (**C and C1**) in the hippocampus of Eu and Ts66Yah mice at different ages 3 months (Eu n = 12, Ts66Yah n = 12); 6 months (Eu n = 12, Ts66Yah n = 12); 9 months (Eu n = 12, Ts66Yah n = 12). Protein levels were normalized per total protein load. All densitometric values are given as percentage of Eu at 3 months set as 100%. Data are presented as means ± SEM. Columns were used to show differences among the groups (Eu vs Ts66Yah). Lines were used to show age-associated changes within each group. For columns: *p < 0.05, **p < 0.01, ***p < 0.001 and ****p < 0.0001. For lines: #p < 0.05, ##p < 0.01, ###p < 0.001 and ####p < 0.0001 among 3 months vs 6 and 9 months; +p < 0.05, ++p < 0.01, +++p < 0.001 and ++++p < 0.0001 among 6 months vs 9 months.

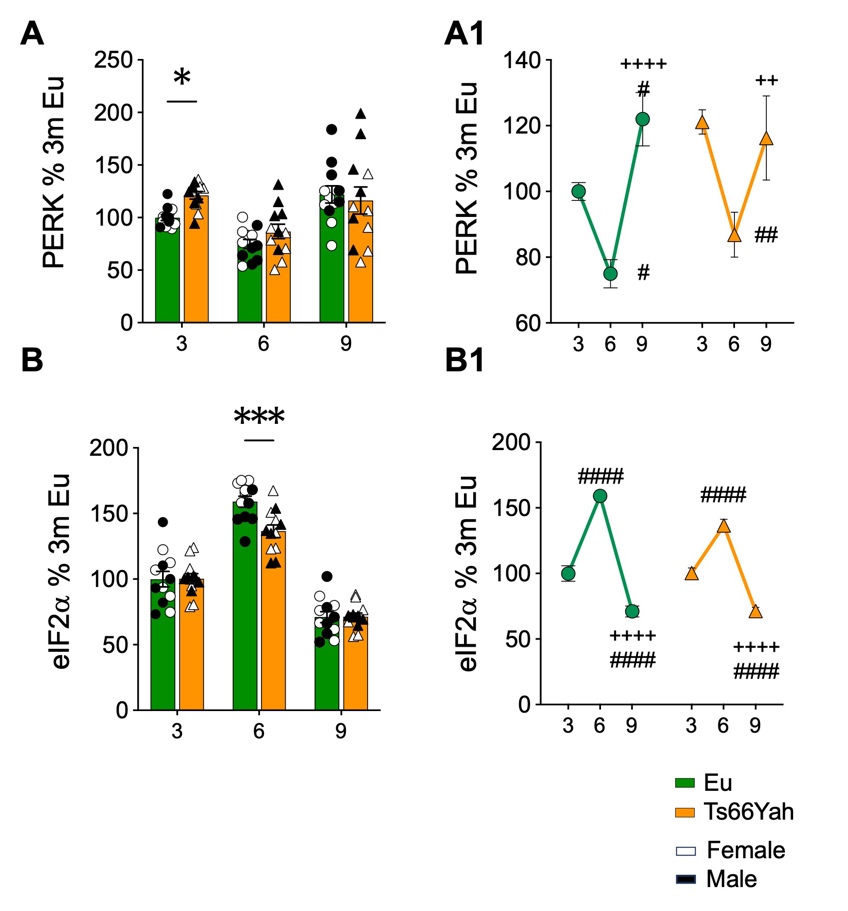

**Supplementary figure 5.** Densitometric evaluation of PERK (**A and A1**) and eif2a (**B and B1**) in the hippocampus of Eu and Ts66Yah mice at different ages 3 months (Eu n = 12, Ts66Yah n = 12); 6 months (Eu n = 12, Ts66Yah n = 12); 9 months (Eu n = 12, Ts66Yah n = 12). Protein levels were normalized per total protein load. All densitometric values are given as percentage of Eu at 3 months set as 100%. Data are presented as means ± SEM. Columns were used to show differences among the groups (Eu vs Ts66Yah). Lines were used to show age-associated changes within each group. For columns: *p < 0.05, **p < 0.01, ***p < 0.001 and ****p < 0.0001. For lines: #p < 0.05, ##p < 0.01, ###p < 0.001 and ####p < 0.0001 among 3 months vs 6 and 9 months; +p < 0.05, ++p < 0.01, +++p < 0.001 and ++++p < 0.0001 among 6 months vs 9 months.
